# Resolving vs. Non-resolving Sphingolipid Dynamics During Macrophage Activation: A Time-resolved Metabolic Analysis

**DOI:** 10.1101/2024.12.31.630925

**Authors:** Nathan F. Chiappa, Nidhi Lal, Edward A. Botchwey

## Abstract

Sphingolipids are increasingly recognized as critical regulators of inflammation and cell fate decisions, with metabolites such as ceramide and sphingosine 1□phosphate exerting contrasting effects on cell survival and proliferation. In macrophages, this balance is especially important, given their central role in host defense, pathogenesis and wound healing. Here, we present a time□resolved model of sphingolipid metabolism in RAW 264.7 macrophages stimulated with KdO□–Lipid A. By integrating measured metabolite concentrations with dynamic flux estimation and established enzyme kinetics, we systematically map dynamic changes in the sphingolipid network during inflammation. Our results reveal a three□phase pattern of sphingolipid remodeling that correlates with distinct functional states of the cell. Moreover, metabolites can be classified into “resolving” or “non□resolving” lipids based on whether they return to basal levels or remain dysregulated through the later phases of the inflammatory response. This partitioning suggests that targeted modulation of specific metabolic nodes may influence the resolution of inflammation. Importantly, our computational approach can assist in the rational design of experimental studies by pinpointing putative drug targets with maximal impact on sphingolipid homeostasis. Such targeted interventions may prevent the pathological amplification of inflammatory signals without globally suppressing essential sphingolipid functions. These findings highlight the utility of an integrative systems□level analysis for elucidating sphingolipid dynamics in macrophages and underscore its potential to guide therapeutic strategies against conditions involving dysregulated inflammation.

## Introduction

Macrophages are key orchestrators of immune responses, functioning at the crossroads of innate and adaptive immunity [1]. They are essential for mounting potent inflammatory responses – producing cytokines, phagocytosing pathogens, and modulating other immune cells – while also contributing to wound healing and tissue homeostasis. However, excessive or dysregulated macrophage activation can drive the pathogenesis of numerous conditions, including atherosclerosis[2], tissue fibrosis [3-5], and cancer [6]. Increasingly, research points to lipid metabolism as a decisive factor in these activation states [7, 8]. Lipids not only supply membrane components for proliferating or migrating cells but also regulate signal transduction pathways that govern macrophage polarization and function. These diverse functional states require precise metabolic coordination, particularly in lipid metabolism, which provides both energy and signaling molecules.

Among lipids, sphingolipids have garnered particular attention for their potent bioactive properties in controlling cell fate and inflammatory processes [6, 9]. Two key metabolites, ceramide and sphingosine-1-phosphate (S1P), often exhibit opposing functions. Ceramide typically favors pro-apoptotic outcomes and cell cycle arrest, whereas S1P supports pro-survival and proliferative outcomes [10-12]. These metabolites are interconnected within a complex metabolic network, where ceramide is created de-novo as well as by the interconversion of complex sphingolipids, and can be converted into sphingosine and subsequently phosphorylated to form S1P [7, 9]. This “sphingolipid rheostat” maintains a delicate balance between pro-survival and pro-apoptotic signals [13], and dysregulation at any point can lead to pathological outcomes like chronic inflammation and tissue damage [3]. Indeed, aberrant sphingolipid metabolism has been implicated in a range of inflammatory pathologies, from metabolic disorders to fibrosis and neurodegenerative conditions [14, 15].

To capture this complexity and identify critical nodes within this network, integrative systems-level approaches are essential [16, 17]. Key enzymes in this network include sphingomyelinases that generate ceramide, ceramidases that convert ceramide to sphingosine, and sphingosine kinases that produce S1P [18]. By merging experimental measurements of metabolite levels with computational modeling, these methods clarify how ceramide, S1P, and their interconversions influence macrophage polarization (e.g., M1 vs. M2 phenotypes), cytokine production, and antigen-presentation capacities [15, 19]. For example, inhibiting acid ceramidase disrupts MHC class II antigen presentation [19], while reinforcing the Spns2/S1P axis prevents hyperinflammation and late-stage immunosuppression [15]. Understanding the timing of these metabolic shifts is crucial for elucidating both protective and pathogenic immune responses [7, 20].

Despite significant progress in mapping sphingolipid pathways, critical gaps remain in our understanding of their temporal dynamics and flux regulation. Most studies measure lipid levels at only a single or limited number of time points [8, 9], providing a static snapshot rather than capturing the dynamic trajectory of how cells transition through different activation states. Furthermore, while large-scale lipidomic databases (e.g., LIPID MAPS) have facilitated the identification and quantification of numerous sphingolipid species [20], few efforts have integrated these data into flux or enzyme-activity analyses to reveal which enzymatic steps truly govern the observed changes [6, 17]. Without this flux information, it remains unclear whether targeting a particular enzyme is a viable anti-inflammatory strategy or would merely induce inconsequential shifts in metabolite levels.

To address these gaps in understanding sphingolipid dynamics during inflammation, we developed an integrated experimental and computational approach combining time-resolved lipidomics with flux balance analysis. This strategy allowed us to address two fundamental questions: 1) How do changes in metabolic flux and enzyme activity drive the temporal evolution of sphingolipid concentrations during macrophage activation? and 2) Can systematic analysis of these fluxes identify optimal targets for anti-inflammatory intervention? By mapping the dynamic relationships between enzyme activities, metabolic fluxes, and lipid concentrations, our work provides a quantitative framework for understanding how macrophages coordinate sphingolipid metabolism during inflammation and reveals new opportunities for therapeutic modulation of inflammatory responses.

## Materials and Methods

### Experimental Design and Data Analysis

Sphingolipids form an interconnected network centered on ceramide, with each metabolite influencing distinct aspects of macrophage function (**Figure 1A)**. To investigate temporal dynamics of sphingolipid metabolism during macrophage activation, we analyzed comprehensive lipidomic data from RAW 264.7 macrophages responding to KdO_2_-Lipid A (KLA) stimulation. Specifically, we utilized the time□course dataset from Dennis et al.[20-23], in which RAW 264.7 macrophages cultured in 10% serum were treated with 100 ng/mL KLA. KLA is a component of lipopolysaccharide (LPS) with confirmed TLR4□mediated bioactivity comparable to LPS [21]. Samples were harvested at 0, 0.5, 1, 2, 4, 8, 12, and 24 hours for both stimulated and unstimulated controls, and LC–MS measurements were collected for a variety of sphingolipids (**Figure 1B)**. For each sphingolipid class, the concentrations of all molecular species (i.e., different acyl□chain variants) were summed at each time point to yield total “pool” concentrations. Because raw measurements in the LIPID MAPS database were provided in units different from enzyme activity values in literature, we converted them to pmol/mg protein, using the relationship that 3 µg of DNA ≈ 0.25 mg protein [17]. Literature□based activity parameters were aggregated to approximate enzyme kinetics in RAW 264.7 cells (**Table** 4). However, since direct measurements for this specific cell line were scarce, composite enzyme activity values from other white blood cell types were used as proxies.

**Figure 1.**
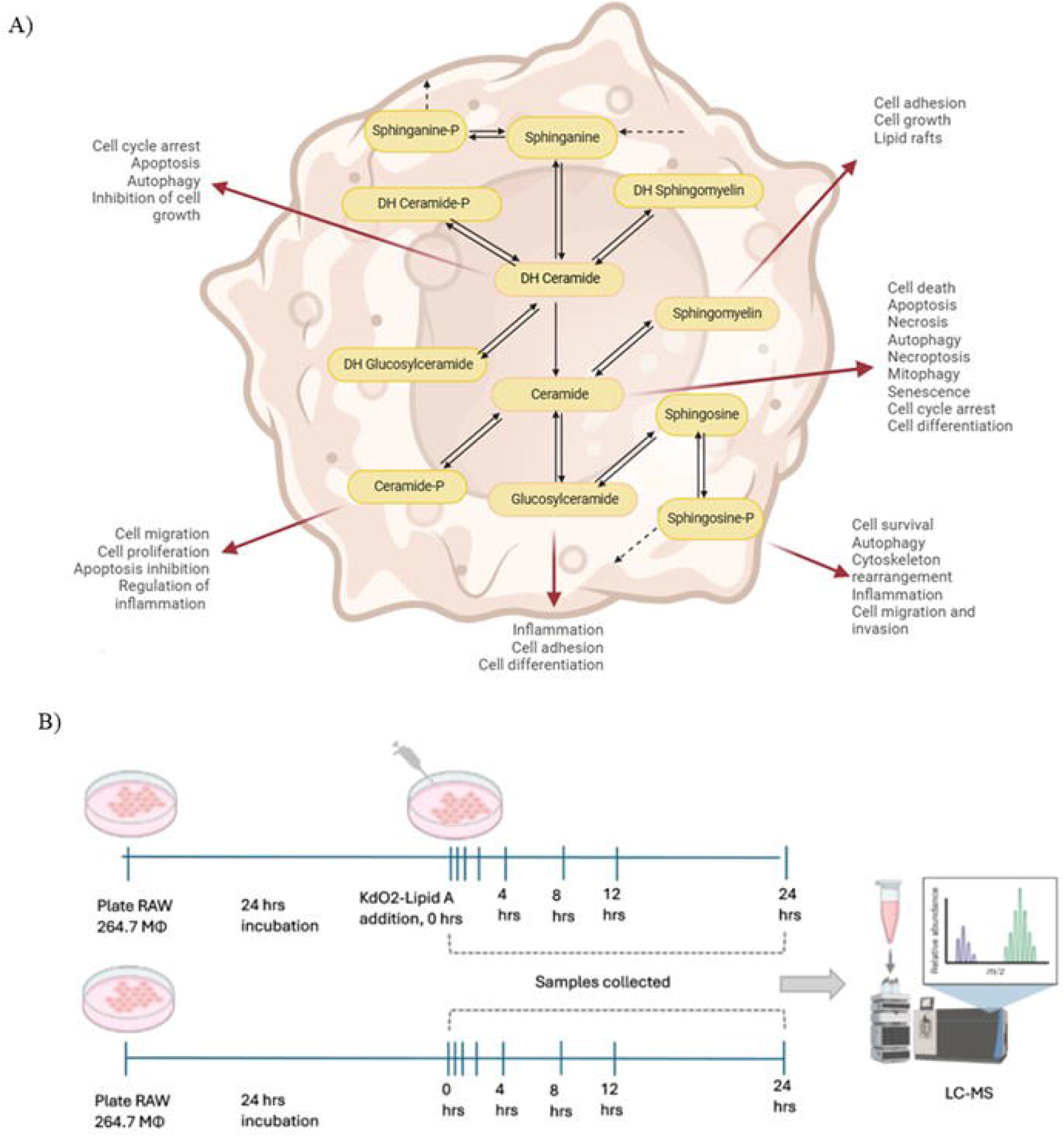
(A) Macrophage sphingolipids as key players in inflammation. This schematic highlights the interconnected sphingolipid network, centered on ceramide as a central “hub” metabolite that links upstream precursors (dihydrosphingosine, dihydrosphingosine□P, dihydroceramide, and their phosphorylated or glycosylated derivatives) to downstream effectors (sphingomyelin, glucosylceramide, sphingosine, and sphingosine□P). Solid arrows denote enzymatic conversions; dashed arrows represent irreversible reactions. Each metabolite exerts distinct yet overlapping roles in regulating macrophage biology—including inflammation, apoptosis, necroptosis, autophagy, cell migration, and survival—thus underscoring the duality of sphingolipid signaling. Ceramide predominantly drives pro□apoptotic and pro□inflammatory responses, whereas sphingosine□P promotes cell survival and cytoskeletal rearrangements. By modulating cell adhesion, proliferation, and differentiation, changes in sphingolipid metabolism can profoundly influence macrophage□mediated inflammation and immune function. **(B)** Experimental design for LIPID MAPS time□course analysis of RAW 264.7 macrophages with or without KdO□□Lipid A stimulation. After 24□hours of initial culture, KdO□□Lipid A (stimulated condition) or vehicle only (unstimulated control) was added, and cells were harvested at 0, 0.5, 1, 2, 4, 8, 12, and 24□hours. Collected samples were processed and analyzed by LC–MS to evaluate temporal changes in the macrophage lipidome.

### Metabolic Network Construction

A sphingolipid metabolic network was constructed based on published literature, the KEGG database, and previously established knowledge of enzyme□mediated reactions. To systematically analyze the temporal evolution of sphingolipid metabolism, we constructed a detailed metabolic network incorporating all measured species and their enzymatic interconversions (**Figure 2A**). The network comprises twelve sphingolipid metabolites (**Table 1**), each assigned a unique variable (X_i_), connected by enzyme-mediated reactions and export processes designated by specific flux numbers (v_i_). Fifteen key enzymes (**Table 2**) catalyze twenty-eight distinct reactions, with twelve additional export processes (**Table 3**). This network structure enabled quantitative flux analysis using the computational workflow (**Figure 2B-C)**.

**Table 1.**
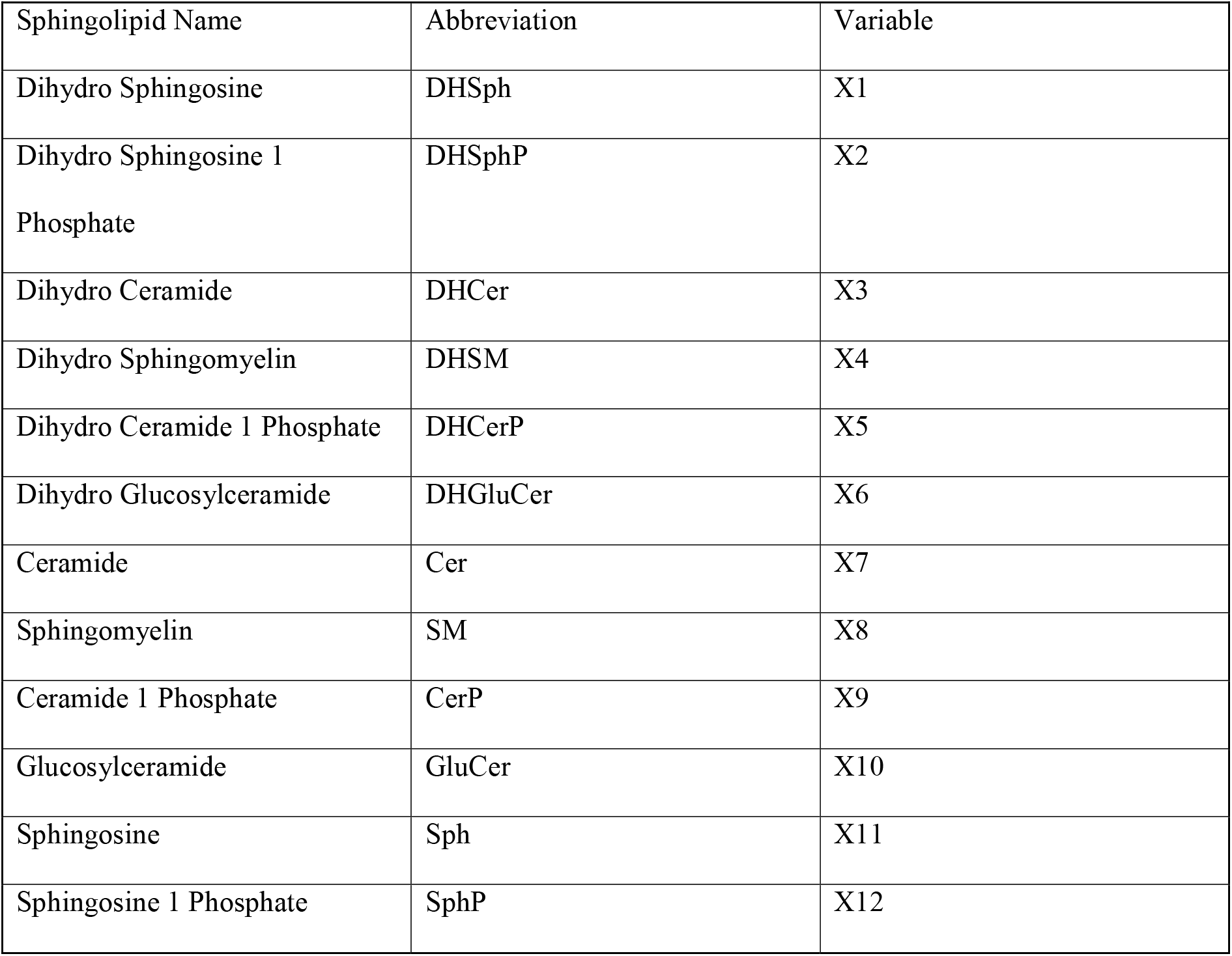
Dependent Variable Identification

| Sphingolipid Name | Abbreviation | Variable |
| --- | --- | --- |
| Dihydro Sphingosine | DHSph | X1 |
| Dihydro Sphingosine 1 Phosphate | DHSphP | X2 |
| Dihydro Ceramide | DHCer | X3 |
| Dihydro Sphingomyelin | DHSM | X4 |
| Dihydro Ceramide 1 Phosphate | DHCerP | X5 |
| Dihydro Glucosylceramide | DHGluCer | X6 |
| Ceramide | Cer | X7 |
| Sphingomyelin | SM | X8 |
| Ceramide 1 Phosphate | CerP | X9 |
| Glucosylceramide | GluCer | X10 |
| Sphingosine | Sph | X11 |
| Sphingosine 1 Phosphate | SphP | X12 |

**Table 2.**
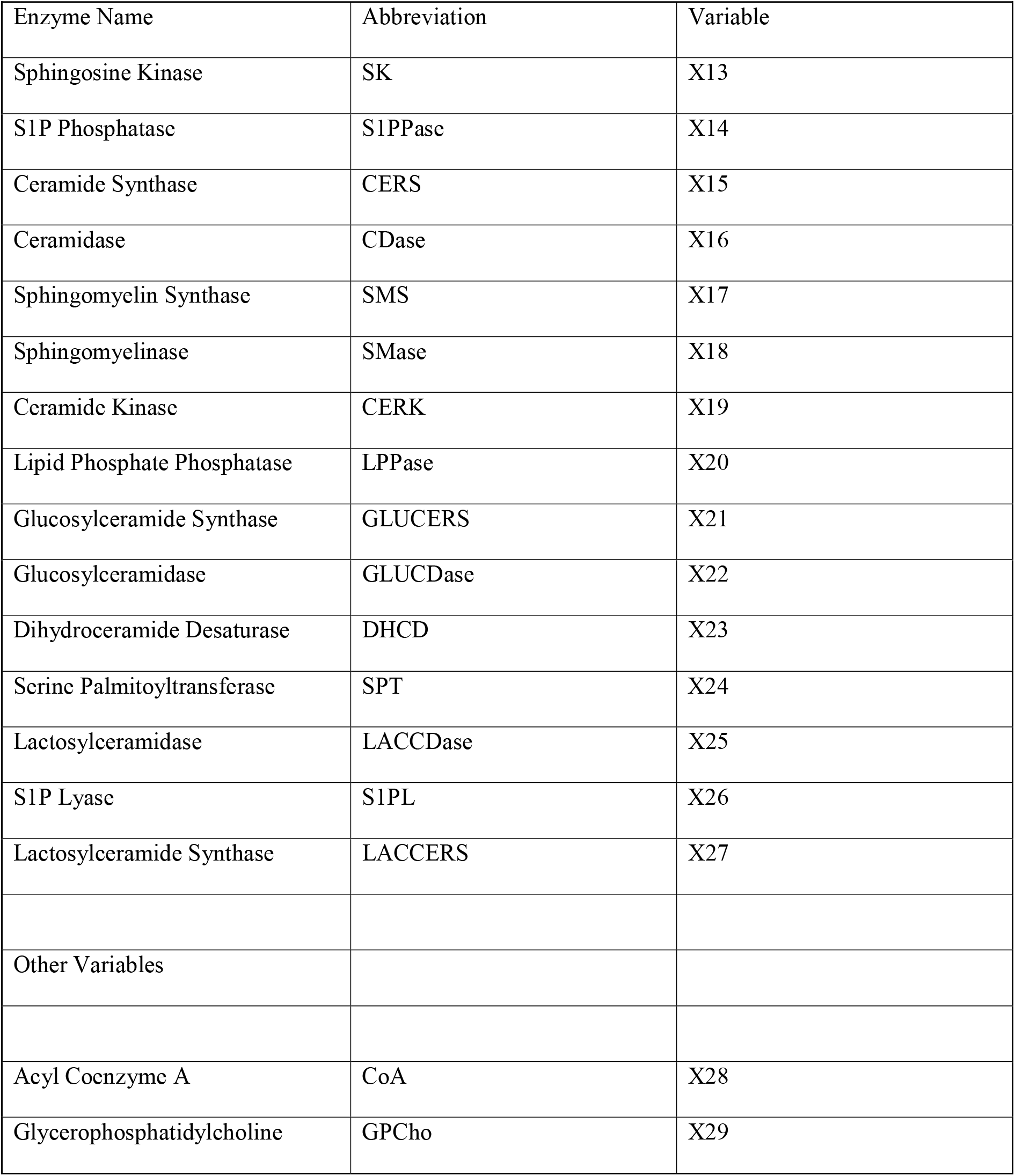
Independent Variable Identification

| Enzyme Name | Abbreviation | Variable |
| --- | --- | --- |
| Sphingosine Kinase | SK | X13 |
| S1P Phosphatase | S1PPase | X14 |
| Ceramide Synthase | CERS | X15 |
| Ceramidase | CDase | X16 |
| Sphingomyelin Synthase | SMS | X17 |
| Sphingomyelinase | SMase | X18 |
| Ceramide Kinase | CERK | X19 |
| Lipid Phosphate Phosphatase | LPPase | X20 |
| Glucosylceramide Synthase | GLUCERS | X21 |
| Glucosylceramidase | GLUCDase | X22 |
| Dihydroceramide Desaturase | DHCD | X23 |
| Serine Palmitoyltransferase | SPT | X24 |
| Lactosylceramidase | LACCDase | X25 |
| S1P Lyase | S1PL | X26 |
| Lactosylceramide Synthase | LACCERS | X27 |
| Other Variables |  |  |
| Acyl Coenzyme A | CoA | X28 |
| Glycerophosphatidylcholine | GPCho | X29 |

**Table 3.**
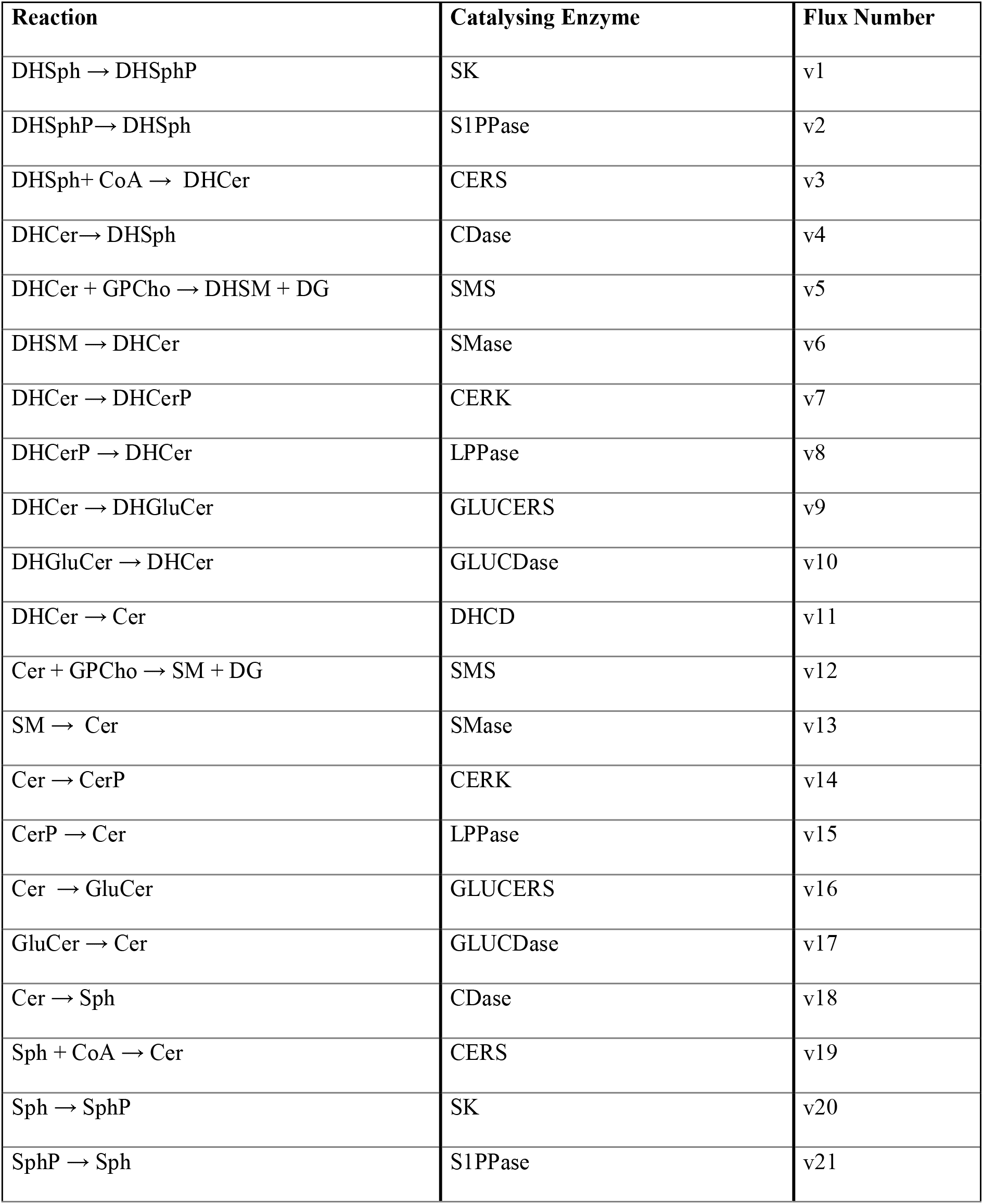
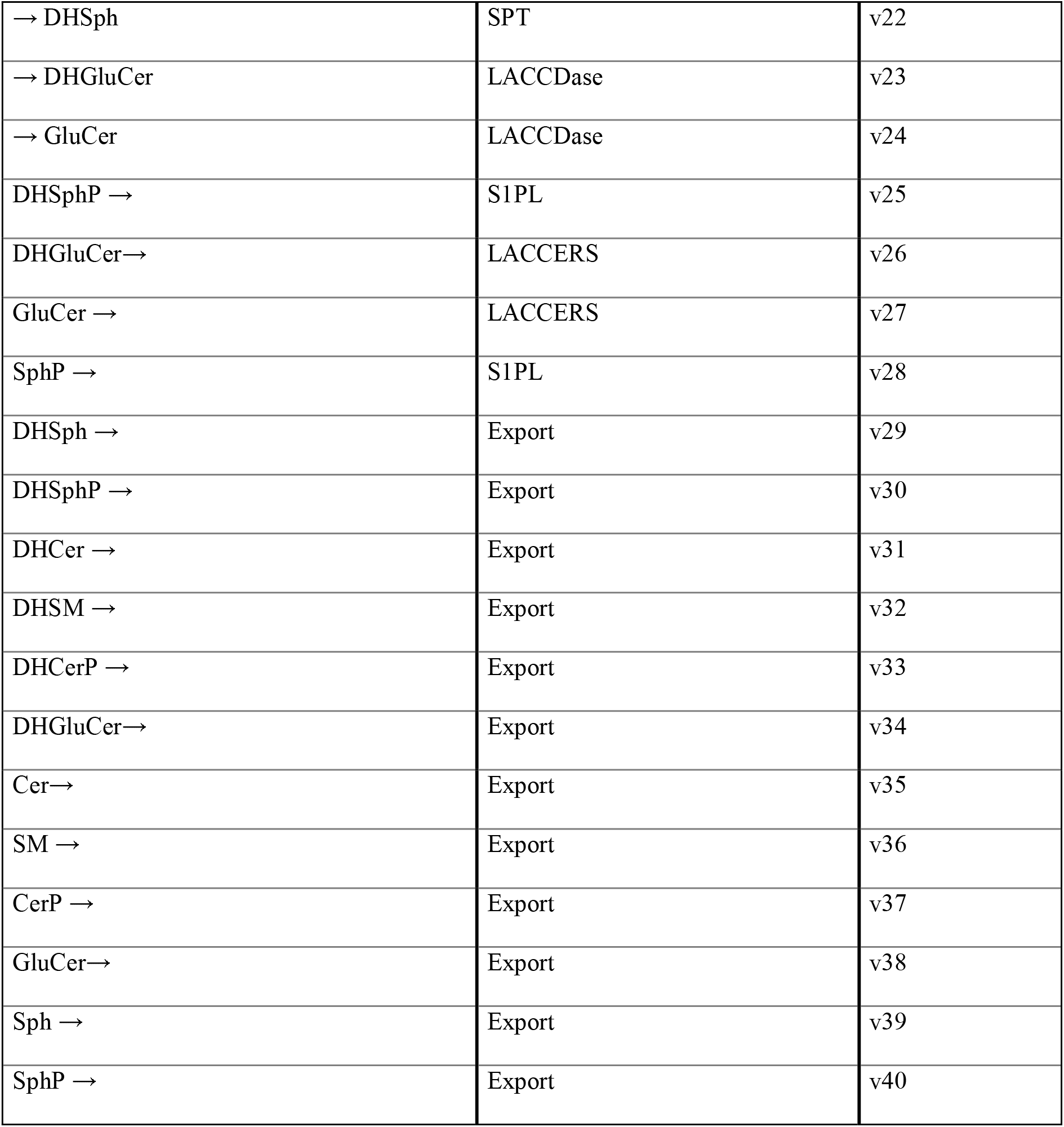
Flux Identification

**Figure 2.**
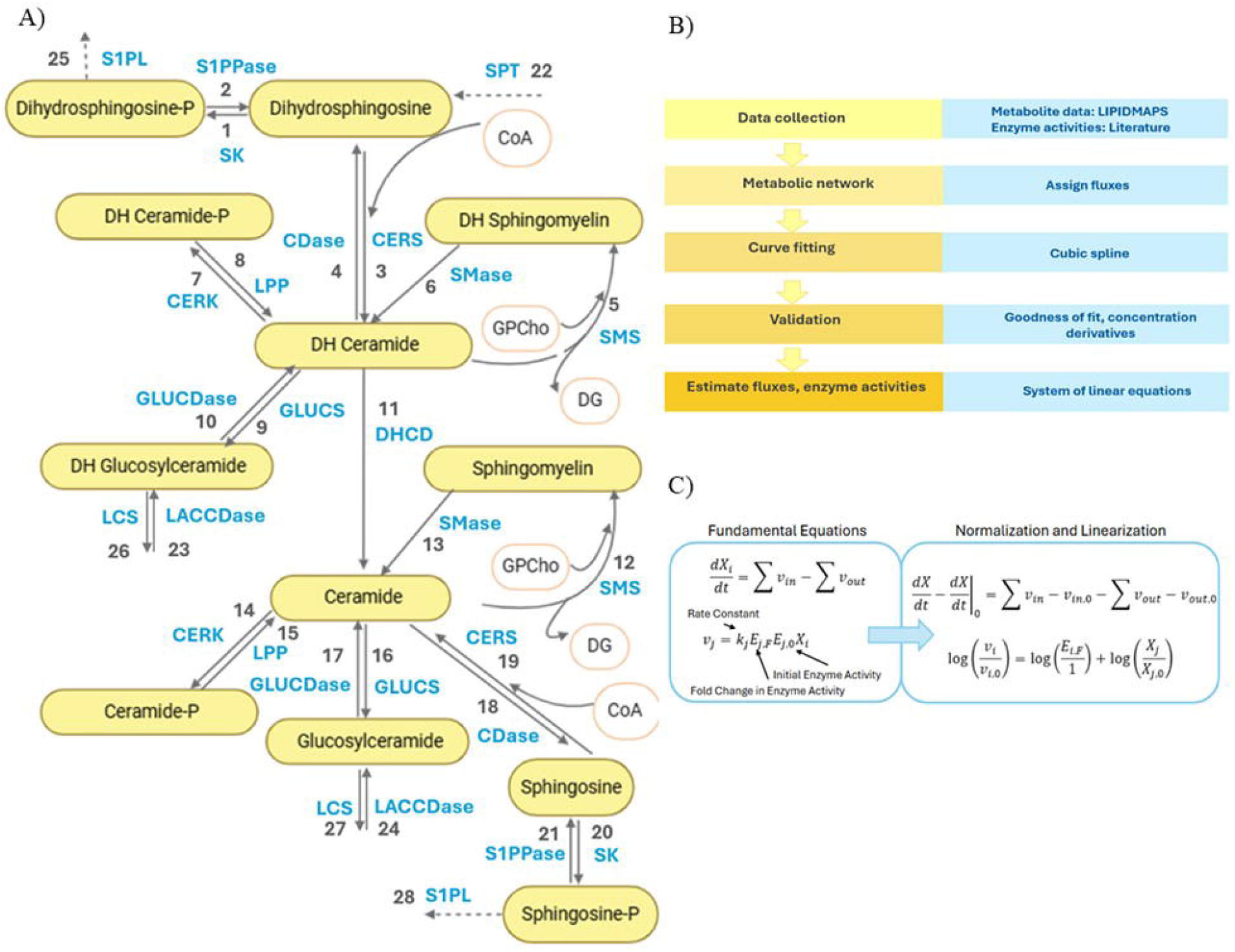
Dynamic flux estimation for the sphingolipid metabolic network. **(A)** Metabolic network diagram showing key enzyme□mediated steps (numbered) in the sphingolipid pathway of KdO□□Lipid A–stimulated RAW□264.7 macrophages. Export processes for each sphingolipid are not shown on the network map. **(B)** Algorithmic workflow for flux estimation. Metabolite concentration data (from LIPID MAPS) and literature□based enzyme activities are integrated into a time□dependent metabolic network. Fluxes are assigned, and a cubic spline–based approach is used for curve fitting. The model is validated by comparing the goodness□of□fit and time□derivatives of measured concentrations, culminating in quantitative estimates of fluxes and enzyme activities. **(C)** Fundamental equations and key variables. Shown are the ordinary differential equations (ODEs) governing metabolite concentrations, along with linearized expressions. Rate constants, initial enzyme activities, and fold changes in enzyme activity are incorporated, enabling dynamic flux inferences from experimental data. Abbreviations: CDase, ceramidase; CERK, ceramide kinase; CERS, ceramide synthase; DHCD, dihydroceramide desaturase; GLUCDase, glucosylceramidase; GLUCS, glucosylceramide synthase; LACCDase, lactosylceramidase; LCS, lactosylceramide synthase; LPP, lipid phosphate phosphatase; S1PL, sphingosine 1□phosphate lyase; S1PPase, sphingosine 1□phosphate phosphatase; SMase, sphingomyelinase; SMS, sphingomyelin synthase; SK, sphingosine kinase; SPT, serine palmitoyltransferase.

### Mass Balance and Rate Equations

The temporal evolution of sphingolipid concentrations can be described through mass balance equations that account for all production and degradation processes. For each metabolite X_i_, the rate of change is given by:

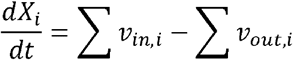

where v_in,i_ represents all fluxes entering X_i_, and v_out,i_ represents all fluxes leaving X_i_. Each flux (v_i_) is associated with an enzymatic (or export) reaction in the network. The general rate equation for a flux v_i_ was:

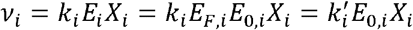

where k_i_ is the rate constant, E_i_ is the effective enzyme activity at a given time, E_0,i_ is the baseline (initial) enzyme activity, E_F,i_ is the fold change of that enzyme relative to baseline, X_i_ is the relevant substrate concentration, and k’_i_ is the product of k_i_ and E_F,i_ which we refer to as the rate parameter. Any unmeasured cofactors were assumed to be in sufficient excess so that their concentration remained effectively constant, and their effects were lumped into the rate constants (k_i_).

**Table 5** illustrates how ceramide (X_7_) participates in multiple reactions (e.g., conversion from dihydroceramide, sphingomyelin, glucosylceramide, and sphingosine). Hence, the ceramide balance equation is: *dX*_7_/*dt* = *v*_11_ + *v*_13_ + *v*_15_ + *v*_17_ + *v*_19_ – (*v*_12_ + *v*_14_ + *v*_16_ + *v*_18_ + *v*_35_) and each flux v_i_ is expressed as: *v*_11_ = *k*_11_*X*_23_*X*_3_, *v*_13_ = *k*_13_*X*_18_*X*_8_, *v*_15_ = *k*_15_*X*_20_*X*_9_, *v*_17_ = *k*_17_*X*_22_***X***_10_, *v*_19_ = *k*_19_*X*_15_*X*_11_*X*_28_ *v*_12_ = *k*_12_*X*_17_*X*_7_*X*_20_, *v*_14_ = *k*_14_*X*_19_*X*_7_, *v*_16_ = *k*_16_ *X*_21_*X*_7_, *v*_18_ = *k*_18_*X*_16_*X*_7_, and *v*_35_ = *k*_35_*X*_7_. Unmeasured substrates (e.g., ATP, UDP-Glucose, or other cofactors) are included in the rate constants k_i_ as they are assumed constant. Combining the flux□balance equations for all sphingolipids yields a vector form: *dX*/*dt* = *Sv* where: X is the vector of sphingolipid concentrations, v is the vector of fluxes, and S is the stoichiometric matrix, specifying how each flux contributes to each sphingolipid’s production or consumption. **Figure 2B** illustrates our overall modeling workflow, and **Figure 2C** summarizes the core rate equations.

**Table 4.**
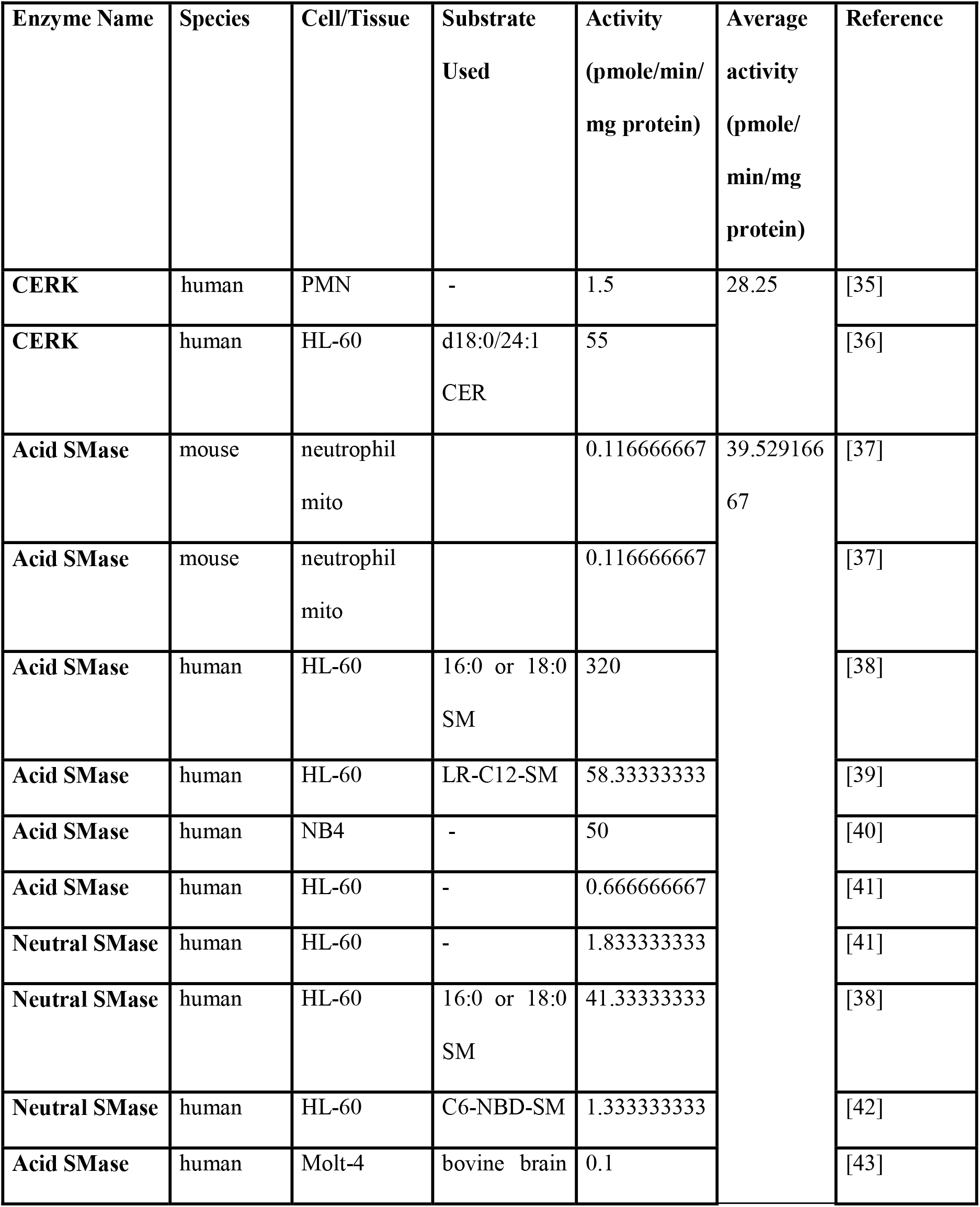
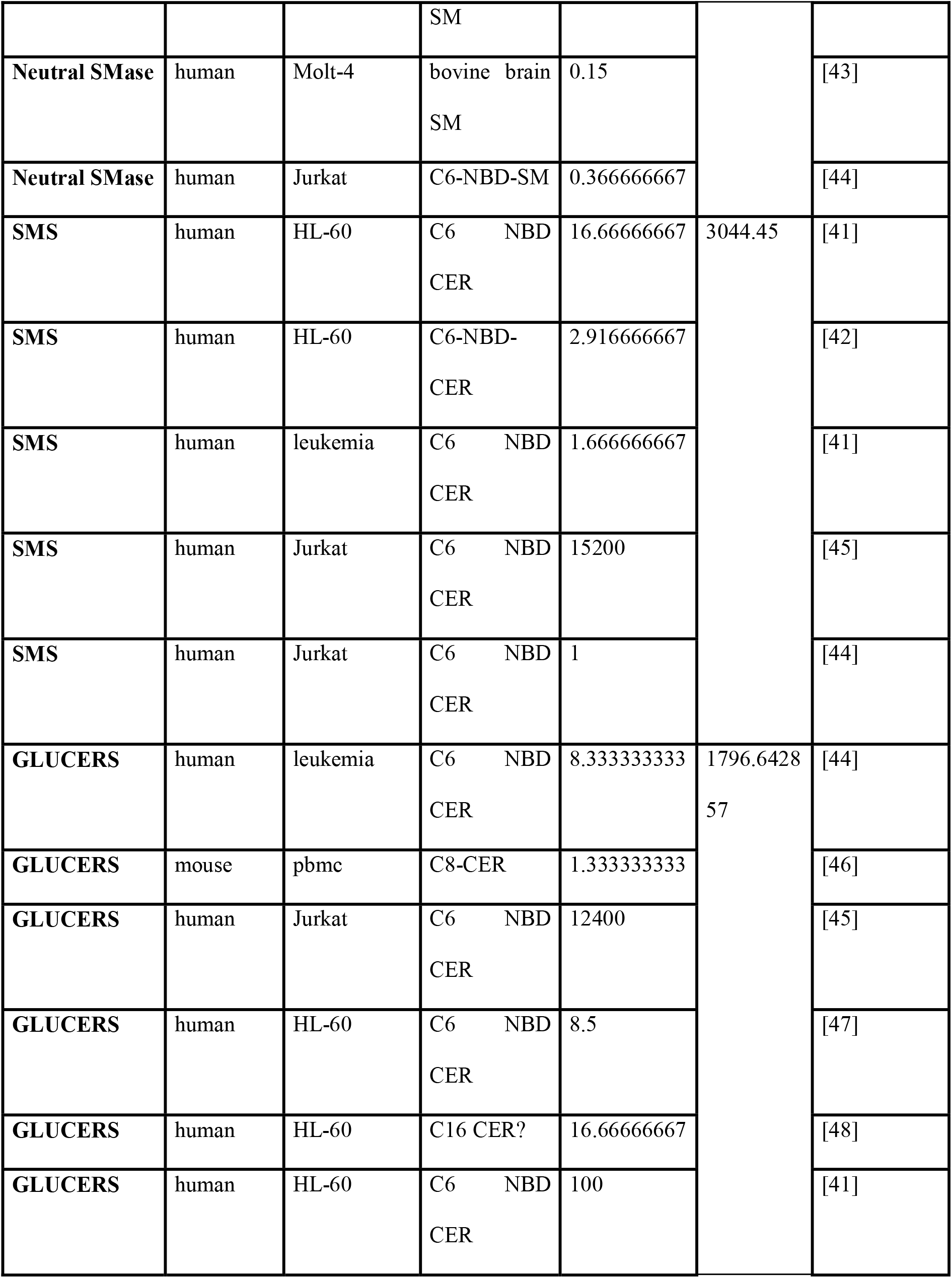
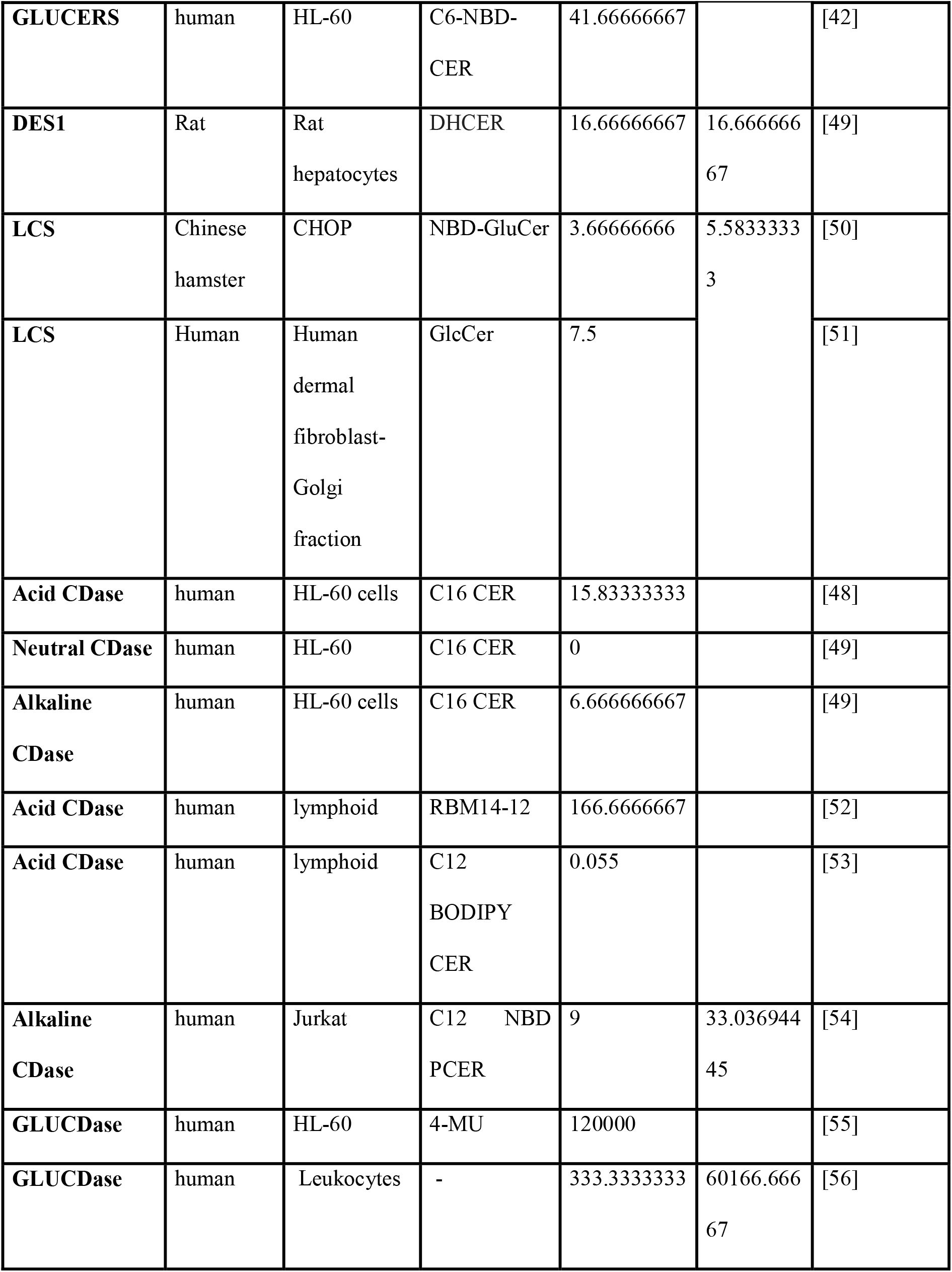
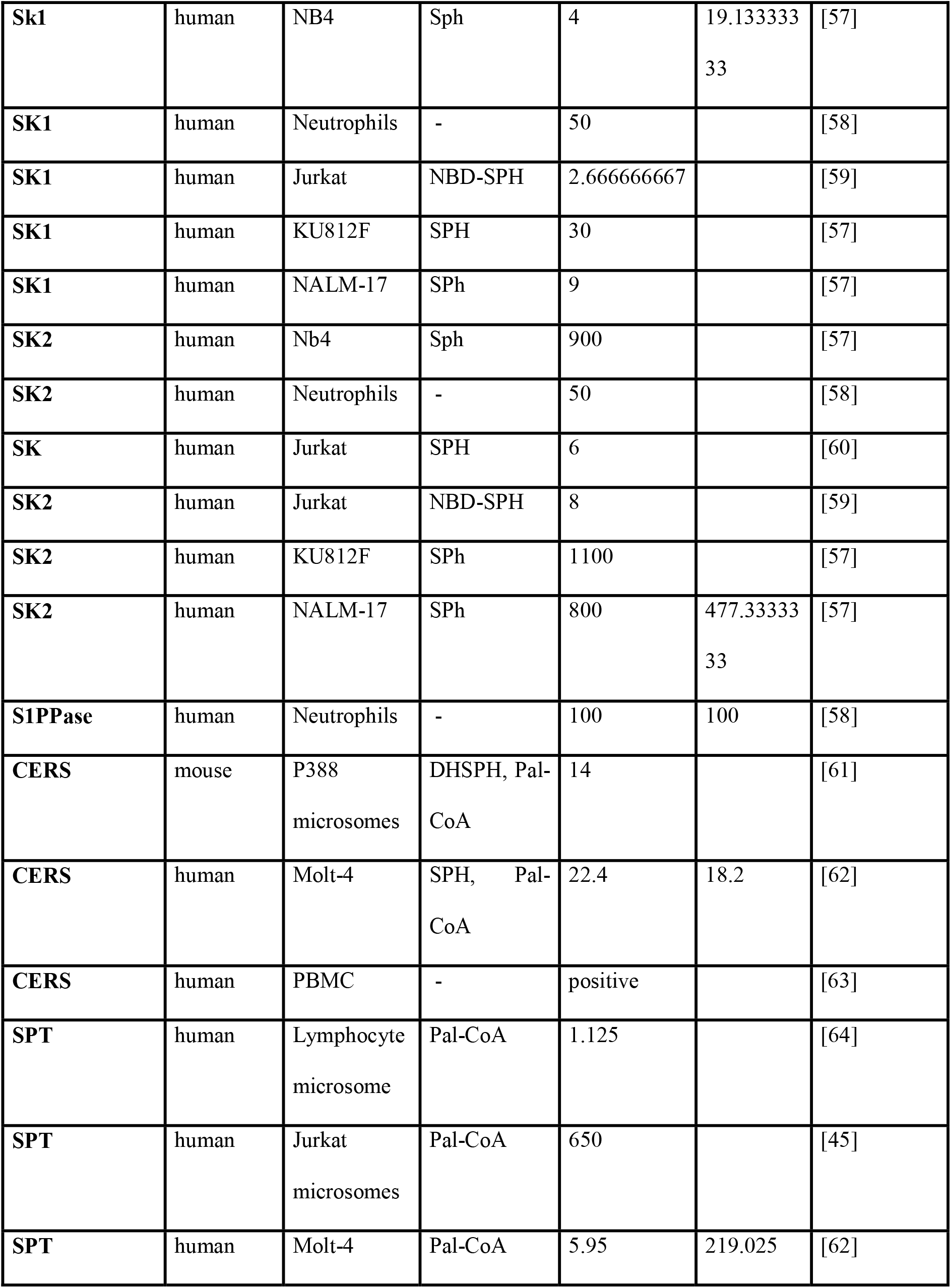
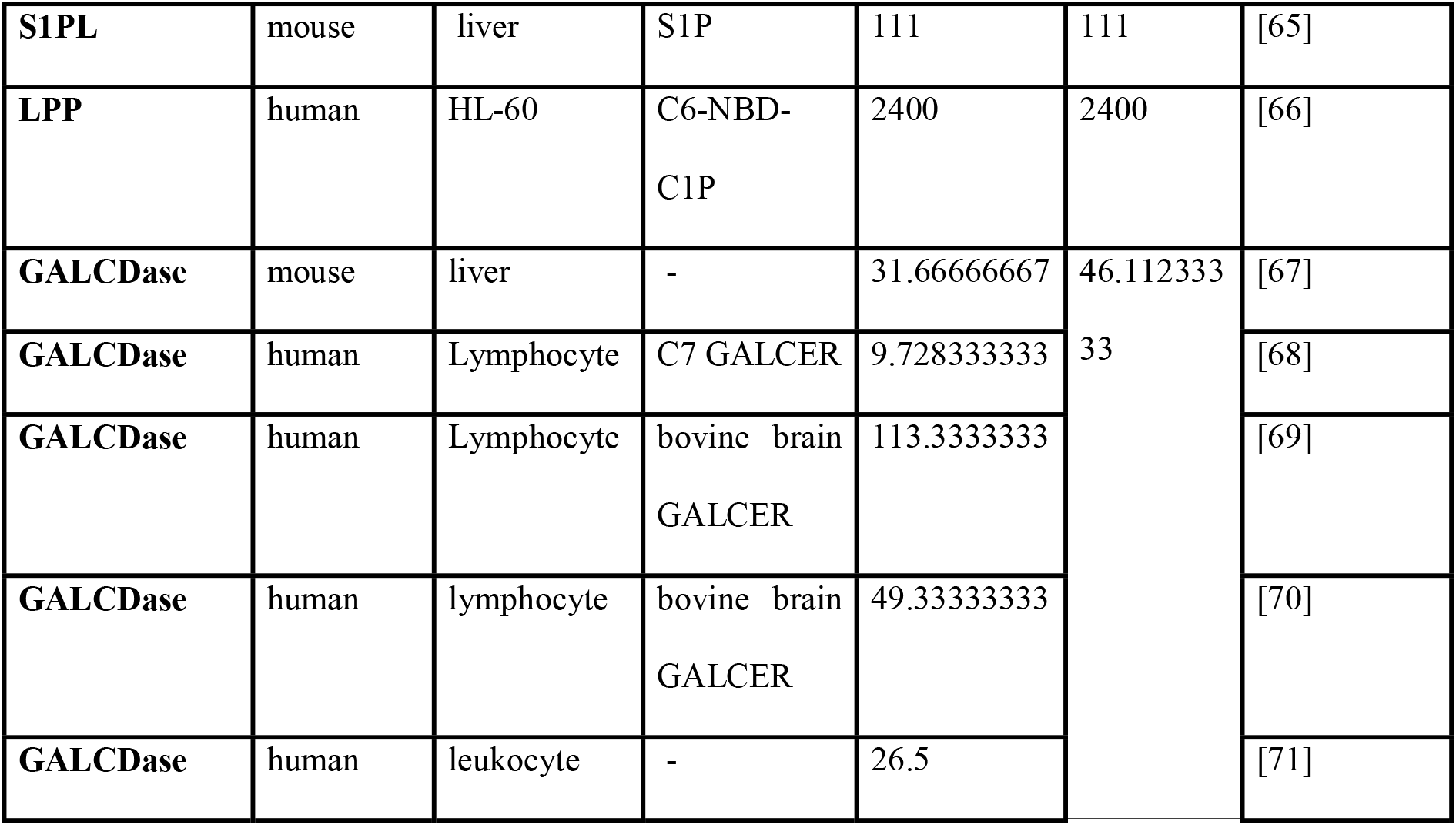
Enzyme Activities Taken from Literature

**Table 5.** Ceramide as an example of flux balance setup

| Flux No. | Reaction | Enzyme |
| --- | --- | --- |
| 11 | Dihydroceramide ($X_3$) → Ceramide ($X_7$) | DHCD ($X_{23}$) |
| 13 | Sphingomyelin ($X_8$) → Ceramide ($X_7$) | SMase ($X_{18}$) |
| 15 | Ceramide 1-phosphate ($X_9$) → Ceramide ($X_7$) | LPP ($X_{20}$) |
| 17 | Glucosylceramide ($X_{10}$) → Ceramide ($X_7$) | GluCDase ($X_{22}$) |
| 19 | Sphingosine ($X_{11}$) + CoA ($X_{28}$) → Ceramide ($X_7$) | CERS ($X_{15}$) |
| 12 | Ceramide ($X_7$) + GPCho ($X_{29}$) → SM ($X_8$) + DG | SMS ($X_{17}$) |
| 14 | Ceramide ($X_7$) → Ceramide 1-phosphate ($X_9$) | CERK ($X_{19}$) |
| 16 | Ceramide ($X_7$) → GluCer ($X_{10}$) | GLUCERS ($X_{21}$) |
| 18 | Ceramide ($X_7$) → Sphingosine ($X_{11}$) | CDase ($X_{16}$) |
| 35 | Ceramide ($X_7$) → Export | — (export) |

### Spline fitting and Derivatives and Numerical Implementation

Metabolic fluxes were estimated over the time course of the experiment using dynamic flux estimation (DFE) [24]. Each sphingolipid’s measured concentration over time was fitted to a cubic smoothing spline (MATLAB’s csaps function) with a manually adjusted smoothing parameter to ensure a visually good fit (**Figure S1**). We then computed time derivatives of these spline fits (using the fnder function), providing numerical values of (dX_i_/dt) at each sampled time point. These derivatives, together with the stoichiometric matrix S, yield a system of equations: *dX*/*dt* = *Sv*. We solved for the flux vector *ν* at each time point by minimizing the sum of squared errors ‖*d***X**/*dt* − *S***v** ‖^2^ via the MATLAB lsqlin function, subject to bounds: *lb* ≤ *v* ≤ *ub*. Lower bounds (lb) for each flux were set to 0 and upper bounds (ub) were set to a multiple of the corresponding enzyme activity taken from literature. The multiple was manually adjusted to ensure the sum of squared errors of the optimal solution was no greater than 10^-10^. From these optimized fluxes, we calculated rate constants and enzyme fold changes. Rate constants (k_i_) were determined from initial conditions: 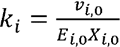 while enzyme fold changes (E_F,i_) were computed throughout the time course: *E*_*F,i*_ = 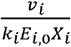. Predicted trajectories for each flux are shown in **Figure S2** and predicted trajectories for each enzyme activity are shown in **Figure S3**.

### Confidence Interval Analysis

The variability present in the LIPID MAPS data set affects the uncertainty in the model predictions. In order to quantify that uncertainty, we took an approach similar to that of Chen et al. [16]. Briefly, for each lipid/time point, we created a normal distribution with the same mean and standard error as the experimental data. We then randomly sampled 1000 points from each normal distribution to create a set of 1000 data sets with the same means and standard errors as the LIPID MAPS data set. We then, performed the same parameter estimation procedure on all 1000 data sets, collected the results and calculated percentiles for each prediction. Since the LIPID MAPS concentration data are shown as mean +/-standard error of the mean, which is the 68% confidence interval of the mean, we show model predictions as the mean and a band containing the middle 68% of the simulation results.

### Global Pattern Analysis

To identify system-wide patterns in sphingolipid metabolism, we performed principal component analysis (PCA) on three key datasets: smoothed metabolite concentrations, calculated fluxes, and inferred enzyme activities. For each analysis, time points were treated as samples, with variables centered and normalized using MATLAB’s pca function. This approach revealed coordinated changes across the sphingolipid network during macrophage activation.

### Dynamic Sensitivity Analysis

The fully parameterized differential equation model enabled investigation of how perturbations in enzyme activities affect sphingolipid concentrations dynamically. To do this, we first define the sensitivity of lipid concentration Xi with respect to enzyme activity Ej as 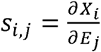. Next, we calculate the time derivative of s as 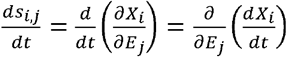. Remembering that 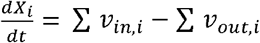, this becomes 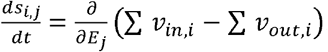. We end up with 12×15=180 differential equations for sensitivities that must be integrated together with the original 12 differential equations for lipid concentrations to generate estimates of dynamic sensitivities [25]. It is often convenient to express sensitivities in relative or scaled terms as 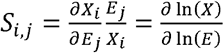. This dimensionless sensitivity coefficient (S_i,j_) measures the relative change in metabolite concentration (X_i_) resulting from a relative change in enzyme activity (E_j_), providing insight into key regulatory points in the network [26].

### Cell Culture

RAW 264.7 cells were expanded in Dulbecco’s Modified Eagle Medium (Gibco 11965118), supplemented with 10% FBS (Corning 35-010-CV) and 1% Penicillin-Streptomycin. The cultures were expanded and seeded onto appropriate well plates at a seeding density of about 105,000 cells/cm^2^ (P4-P5) and allowed to attach overnight. Cells in the treatment group were dosed with 100 ng/ml KdO2-Lipid A (Avanti 699500).

### Flux Tracking

Metabolic fluxes were evaluated using a protocol modified from Nikolova-Karakashian [27]. 10uM C6-NBD SM as a tracer was added at 0 hrs to both control and KLA treated groups. Media and cell (PBS suspension) samples were collected at 0, 4, 12 and 24 hrs after KdO2-Lipid A, where the media samples were centrifuged at 170 x g for 7 minutes to remove any floating cells. Media was collected from the wells and the cells were washed once with PBS and scraped to detach them from the surface. Samples were stored at -80° C and later thawed on ice. They were then sonicated, and 50 uL of each sample was aliquoted for total protein estimation using BCA assay. 400uL of the rest of the samples were aliquoted into glass lipid extraction tubes. C6 NBD DHCer as an internal standard, as well as 1 ml of NBD-Sphingolipid mobile phase consisting of 850 mL MeOH, 150 mL H_2_O and 1.5 mL Phosphoric acid, was added and allowed to incubate in a 48° C water bath overnight with shaking. The samples were centrifuged at 1000xg for 10 minutes, and the top 1mL of the solution was transferred to HPLC autosampler vials without disturbing the bottom to remove settled material. The samples were analysed using a Shimadzu HPLC with fluorescence detector (ex/em: 460/535 nm). Peak areas were converted to concentrations using a calibration curve.

Metabolic fluxes were estimated using a simplified flux balance model including only the NBD-lipid metabolites in the media and in the cells. The differences in fluxes between the KLA-stimulated cells and the control cells were calculated and compared to the model predictions using the LIPID MAPS data. NBD-lipid concentrations with curve fitting are shown in **Figure S6**.

### Enzyme Activity Assay

Acid and neutral SMase activites were evaluated using a protocol modified from Nikolova-Karakashian [27]. Media was removed from the wells and the cells were washed with PBS before scraping in PBS and storing at -80° C. The samples were later thawed on ice and sonicated for cell lysis. 50 uL of each sample was aliquoted for total protein estimation using BCA assay. Based on the assay, an amount of sample containing 5ug of protein was added into Eppendorf tubes and 400uL of the appropriate buffer (acid/neutral) containing 5uM C6-NBD SM was added into each tube. The acid buffer is a 0.5 M acetate buffer containing sodium acetate and glacial acetic acid in DI water, pH 4.5 and the neutral buffer is a 10 mM Tris Buffer containing Tris HCl, Tris base, 5 mM MgCl2 and 0.2% (w/v) Triton X-100, pH 7.4. The reaction tubes were placed in a shaking water bath at 37° C for 1 hour for the reaction to proceed. 1ml of the NBD-Sphingolipid mobile phase described previously was added to each tube. The samples were transferred to lipid extraction tubes and put back in the shaking water bath for another hour. They were then centrifuged at 1000g for 10 minutes to pellet insoluble material. 1ml of the supernatant was transferred to autosampler tubes and analysed by a Shimadzu HPLC with fluorescence detection (ex/em: 460/535 nm).

### Total Protein Estimation

Total protein was estimated to be using a Pierce BCA assay kit (**23225**). 0, 0.05, 0.1, 0.2, 0.4, 0.8 and 2 mg/mL BSA was used as assay standards. 10 uL of the media or cell homogenates was added in triplicate to a 96 well plate and subsequently 190 uL of the BCA reagent solution containing reagents A and B in the ratio 50:1 was added. The plate was covered with aluminum foil and incubated for 30 mins at 37°C and read on a plate reader (SpectraMax M2, Molecular Devices) at 562 nm.

## Results

### PCA Reveals a Three□Phase Sphingolipid Response

Using the LIPID MAPS sphingolipid concentration data (**Figure S1**) and our flux model algorithm, we generated predictions of metabolic fluxes (**Figure S2**) and enzyme activities (**Figure S3**). However, the number of these variables and their dynamic nature make global insights illusive. To gain a global overview of how sphingolipid metabolism evolves over the 24□hour time course, we performed principal component analysis (PCA) on all three datasets—lipid concentrations, metabolic fluxes, and enzyme activities—extracted from KLA□stimulated RAW 264.7 macrophages. Despite measuring different levels of the pathway, all three PCA plots (**Figure 3A–C**) revealed a similar trajectory that naturally separates into three phases: Phase 1 (0–4 h), Phase 2 (4–12 h), and Phase 3 (12–24 h). These phases are indicated by shifts in the first two principal components (PC1 and PC2), which together explain the majority of the variance in each dataset. The color scale in **Figure 3** highlights time progression, and arrows denote the transition boundaries between phases.

**Figure 3.**
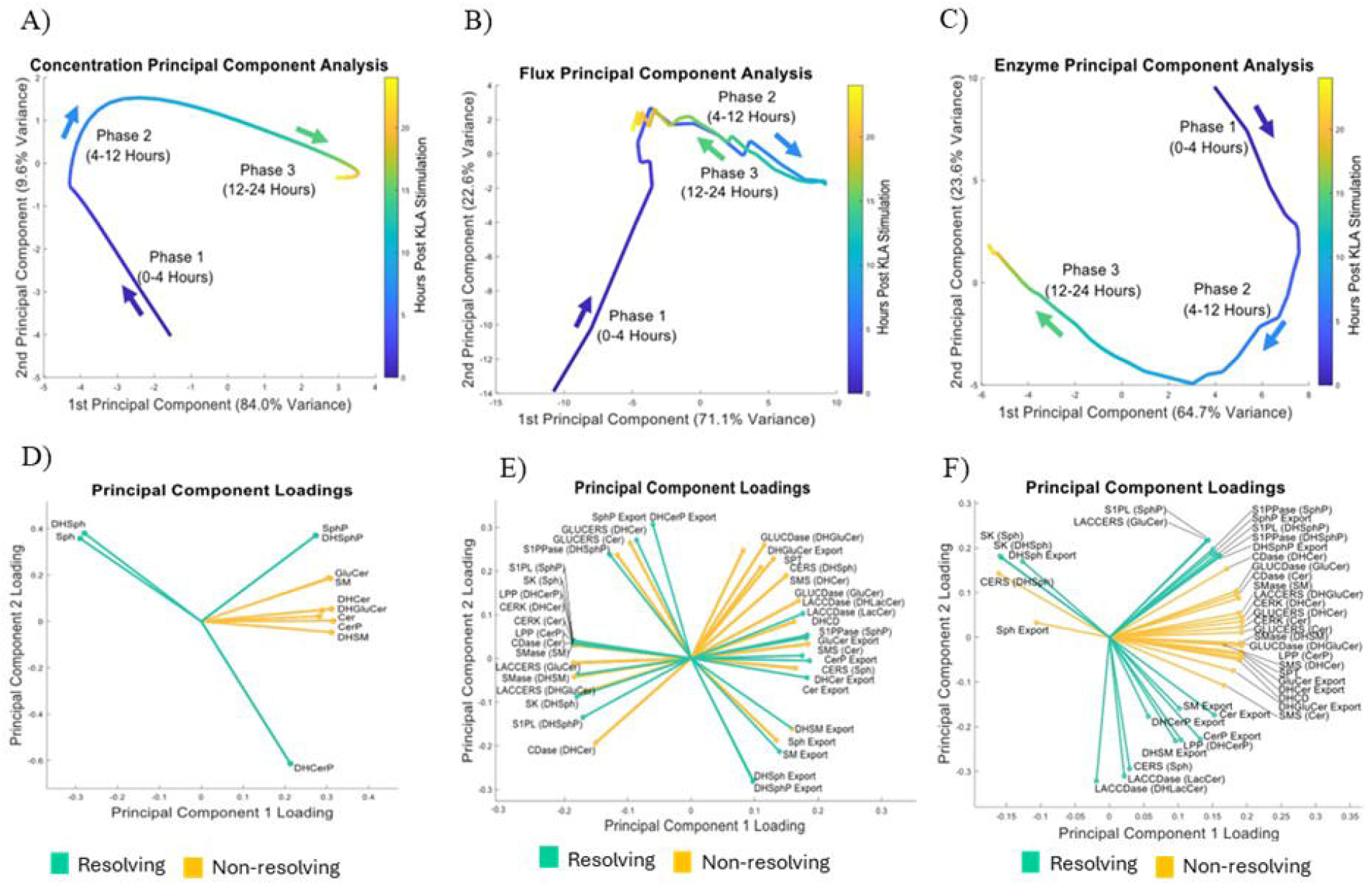
Principal component analysis (PCA) reveals a three□phase response in macrophage sphingolipid metabolism. **(A–C)** PCA of **(A)** concentrations, **(B)** fluxes, and **(C)** enzyme activities each displays a similar three□phase trajectory (Phase□1: 0–4□hours, Phase□2: 4–12□hours, Phase□3: 12–24□hours). The color scale indicates sampling time; arrows highlight transitions between phases. Despite measuring different levels (lipids, fluxes, or enzymes), all datasets shift coherently from early to intermediate to late time points, suggesting coordinated but distinct metabolic dynamics. **(D–F)** Loadings plots for the first two principal components show which sphingolipid species **(D)**, fluxes **(E)**, and enzymes **(F)** most strongly drive each phase of the response. Variables with large positive or negative loadings on PC1 or PC2 are key contributors to overall variance. Clustering patterns help pinpoint specific molecular processes (e.g., ceramide synthesis or sphingosine phosphorylation) that define early or late metabolic transitions. Overall, these PCA results confirm that the macrophage sphingolipid response to KdO□□Lipid A proceeds in three overlapping phases, highlighting how multiple layers of metabolism (concentrations, fluxes, and enzymes) coordinate to shape phase□specific adaptations. Figures E and F show flux and enzyme labels with the corresponding substrates in parentheses, where the same enzyme can metabolize multiple substrates. Since fluxes completely resolve by the end of phase 3, the color coding of fluxes into resolving and non-resolving groups is based on the enzyme that catalyzes that reaction.

Strikingly, each of the three PCA plots exhibits a smooth, coherent transition from early (Phase 1) to intermediate (Phase 2) and then to late (Phase 3) time points, suggesting that sphingolipid concentrations, fluxes, and enzymes are coordinately remodeled as inflammation proceeds. Notably, while the largest changes in PC scores occur between 4 and 12 hours (transition into Phase 2), cells also appear to enter a distinct late□stage metabolic configuration after 12 hours. This late shift underscores the possibility that certain lipids and enzymes remain persistently altered or begin to resolve only by the end of the experiment.

To identify which specific sphingolipid species, fluxes, or enzymes dominate each phase, we examined the loadings plots for PC1 and PC2 (**Figure 3D–F**). Key metabolites such as ceramide, sphingosine, and their phosphorylated forms often show high loadings, indicating they explain a substantial portion of the variance and help define major time□dependent transitions. Similarly, fluxes associated with sphingomyelin synthase (SMS) or ceramidase (CDase), and enzyme activities including sphingosine kinase (SK) and sphingomyelinase (SMase), cluster in ways that differentiate Phase 1 from Phase 2/Phase 3. In these figures, the notation “Enzyme (Substrate)” indicates that the enzyme is capable of using multiple substrates and the enzyme uses the indicated substrate for that reaction. These loadings patterns point to distinct metabolic programs across early, intermediate, and late inflammatory responses—for example, ceramide production vs. sphingosine phosphorylation— and reinforce the concept that macrophages rewire their sphingolipid metabolism in stage□specific patterns. Such phase□wise coordination implies that macrophages do not simply increase or decrease a single sphingolipid species; rather, they dynamically restructure the entire network in ways that likely underlie functionally distinct inflammatory and resolution states.

### Partial vs. Complete Resolution of Sphingolipid Metabolism by Phase 3

To determine the extent of recovery in sphingolipid metabolism over the 24-hour time course, we compared stimulated vs. control cells in terms of (A) sphingolipid concentrations, (B) fluxes, and (C) enzyme activities using scaled principal component (PC) scores (**Figure 4A–C)**. In the sphingolipid and enzyme datasets (**Figure 4A,C**), PC2 largely returns to near□control levels by 24 hours, but PC1 remains markedly shifted. This suggests that, by the end of Phase 3, a subset of lipids and enzymes does revert toward pre□stimulation values (“resolving” components), whereas others remain altered (“nonresolving”) and could potentially drive sustained inflammatory or pro□survival signaling.

**Figure 4.**
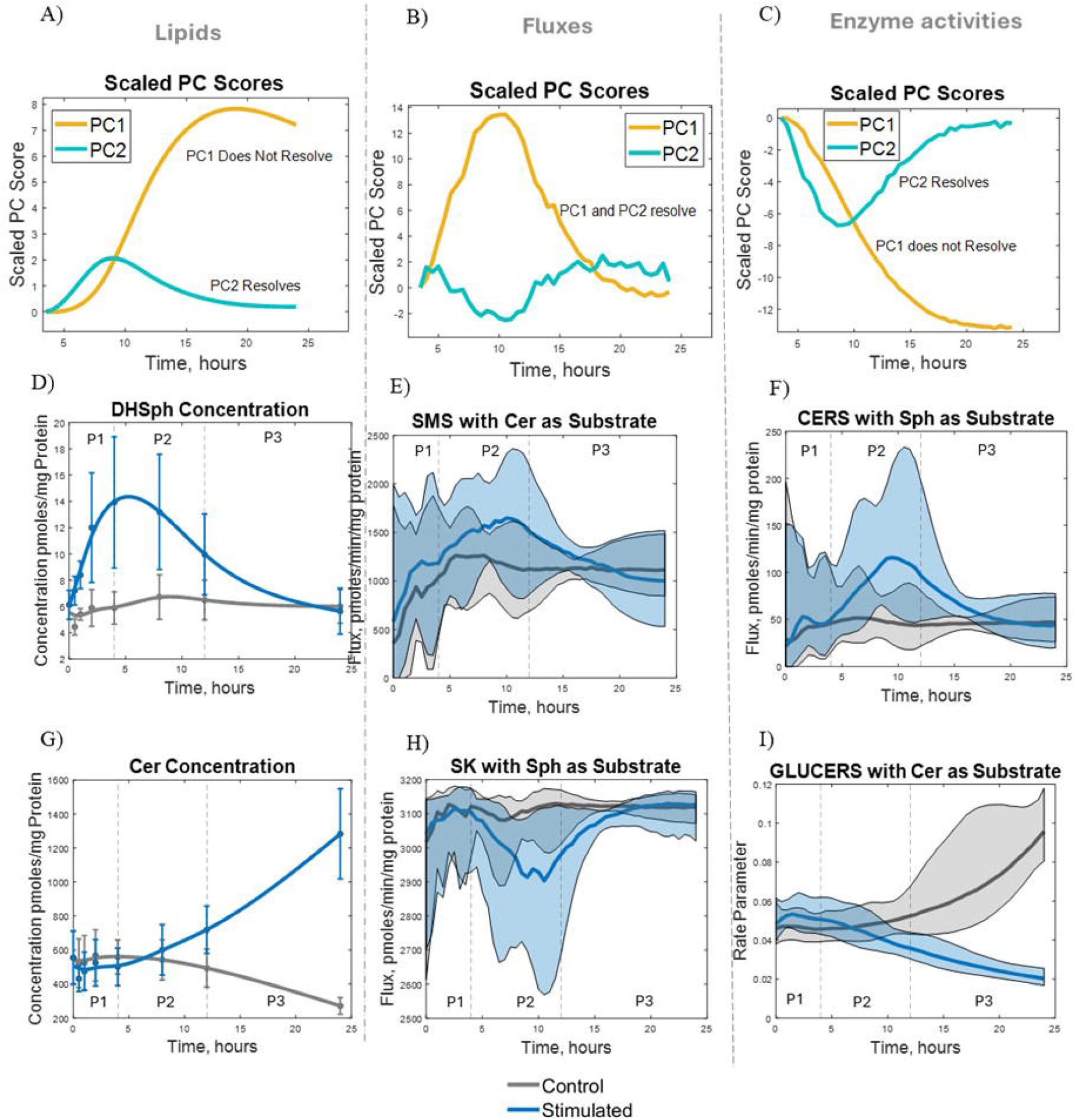
Time□course analyses reveal partial resolution of many sphingolipid species and enzyme activities—but complete resolution of fluxes—by Phase 3. **(A–C)** Scaled principal component (PC) scores over time for sphingolipid concentrations **(A)**, fluxes **(B)**, and enzyme activities **(C)**. PC2 largely returns to baseline by the end of Phase□3 (partial resolution) in the sphingolipid and enzyme datasets, whereas PC1 remains shifted. In contrast, both PC1 and PC2 resolve in the flux dataset by the end of Phase□3. **(D–F)** Representative examples of resolving behaviors for specific lipids **(D)**, fluxes **(E)**, and enzyme activities **(F). (G–I)** Representative examples of non-resolving behaviors for specific lipids **(G)**, fluxes **(H)**, and enzyme activities **(I)**. Each plot shows the time course under stimulated (blue) and control (gray) conditions, with the three phases indicated. “Resolving” variables return near control levels by the end of Phase□3, whereas “nonresolving” ones remain elevated or suppressed. Vertical dividers mark Phase□1 (0–4□h), Phase□2 (4–12□h), and Phase□3 (12–24□h). These examples illustrate how the overall PCA trajectories **(A–C)** are driven by diverse molecular outcomes at the level of individual lipids, fluxes, and enzymes. In (**D-I)**, P1, P2 and P3 refer to phases 1, 2 and 3 respectively. In (**E, F, H**, and **I)**, the colored bands around the mean contain the middle 68% of estimation results.

By contrast, both PC1 and PC2 in the flux dataset (**Figure 4B**) return to baseline levels by 24 hours. Thus, at the level of reaction rates, the macrophage sphingolipid network appears to restore its global flux patterns, even if certain metabolites and enzymes do not completely normalize. This mismatch between flux recovery and partial metabolite/enzyme recovery indicates that, after 12–24 hours, cells maintain a stable flow of sphingolipid intermediates but with altered pool sizes or enzyme expression.

To illustrate these behaviors, **Figure 4D–F** present representative time□course plots of “resolving” concentrations, fluxes, and enzyme activities while **Figure 4G-I** present representative plots of “nonresolving” concentrations, fluxes, and enzyme activities. Vertical lines at 4 hours and 12 hours delineate the same three phases identified earlier—Phase 1 (0–4 h), Phase 2 (4–12 h), and Phase 3 (12–24 h). For instance, one sphingolipid class may return near the control level (**Figure 4D**, resolving) while another remains elevated through 24 hours (**Figure 4G**, nonresolving). A similar dichotomy emerges in enzyme activities (**Figure 4F and I**), showing how certain nodes in the pathway can reset by Phase 3, whereas others appear locked in an inflammatory or adaptive state. These findings underline the multi□layered complexity of macrophage sphingolipid regulation, where dynamic shifts in flux can compensate for persistent deviations in metabolites or enzyme levels to achieve a quasi□homeostatic state by the end of Phase 3. In **Figure 4 E, F, H, and I**, the confidence intervals of the flux and enzyme activitiy predictions are shown as colored bands around the mean predictions.

### Validation of Model Predictions

To validate the model’s predictions of fluxes and enzyme activities using orthoginal methods, we estimated metabolic fluxes using C6-NBD Sphingomyelin as tracer and measured the activity of sphingomyelinase (acid plus neutral) as a representative enzyme. **Figure 5A** and **B** show the experimental design for the two experiments. **Figures 5C-E** show that the difference in the model’s predictions of flux between the control and KLA treated conditions matches with that obtained experimentally using the NBD-sphingolipid tracer, for the fluxes through sphingomyelinase, glucosylceramidase and lactosylceramidase. Similarly, the difference in total SMase activity observed using the NBD-substrate matches with that of the original model predictions (**Figure 5F**).

**Figure 5.**
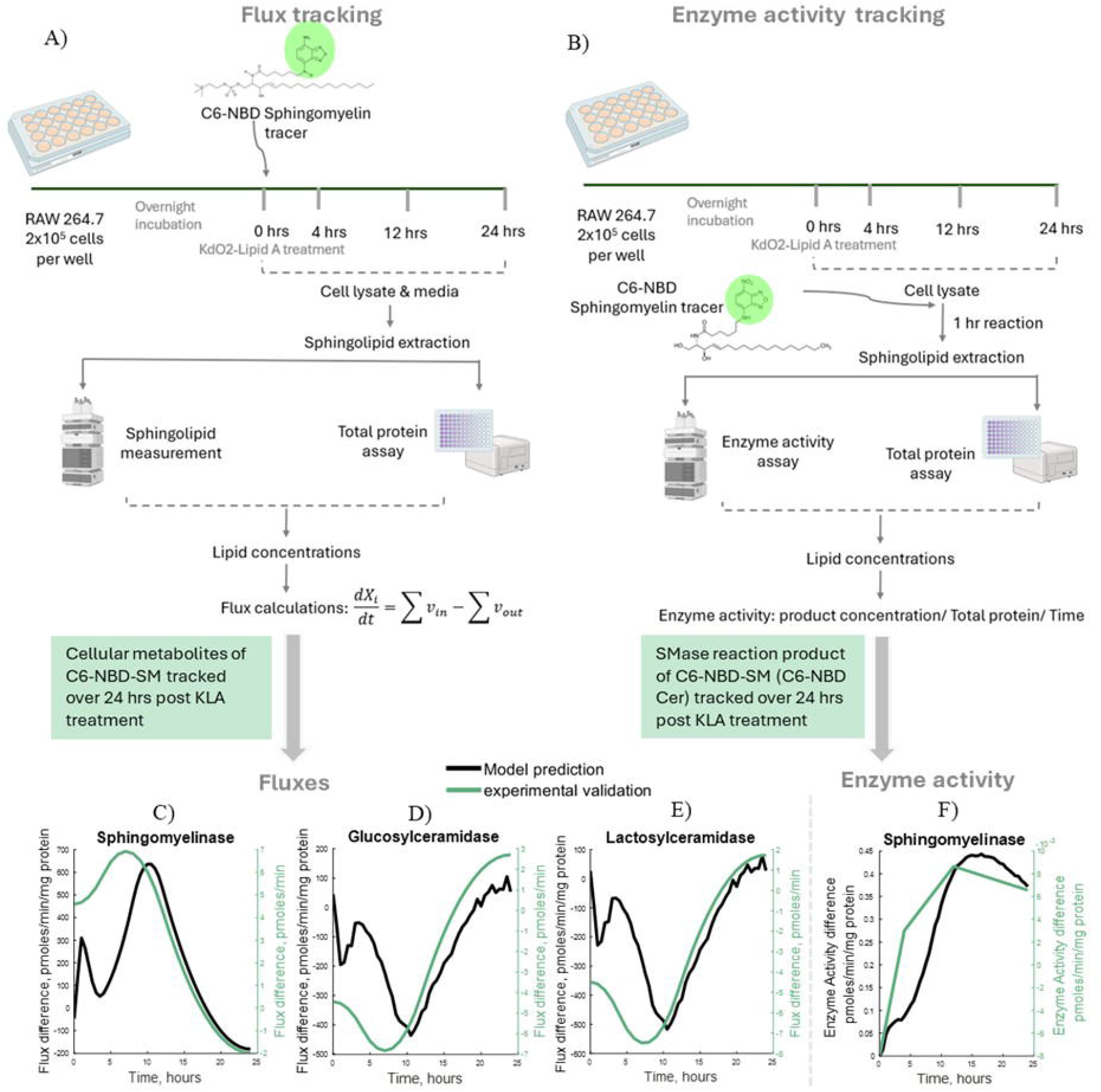
Experimental validation of model predictions. **(A)** Experimental design for estimation of metabolic fluxes using C6-NBD-Sphingomyelin as a tracer. **(B)** Experimental design of sphingomyelinase enzyme activity measurement using C6-NBD-Sphingomyelin as a substrate. **(C-E)** Experimentally determined fluxes trace the same trajectory as the model prediction for **(C)** Sphingomyelinase, **(D)** Glucosylceramidase and Lactosylceramidase **(E)**. Plots are shown as the difference in fluxes between unstimulated and KLA stimulated conditions. **(F)** Experimentally determined enzyme activity of total sphingomyelinase traces the same trajectory as the model prediction. Plot is shown as the difference in activity between unstimulated and KLA stimulated conditions.

### Phase□Specific Shifts in the Sphingolipid Network Reflect Coordinated Flux Modulations

To better visualize how fluxes and concentrations co□evolve across the three phases, we mapped the observed changes onto the sphingolipid network for Phases 1, 2, and 3 (**Figure 6B–D**). Consistent with the flux analyses, Phase 1 (0–4 h) is characterized by a net flow from complex sphingolipids (such as sphingomyelin, ceramide 1 phosphate, and ceramide) toward long□chain bases (sphingosine, dihydrosphingosine). In **Figure 6B**, this manifests as red arrows or boxes for several complex species, indicating decreases in their concentrations/fluxes, paired with green boxes/arrows for Sph and DHSph routes. This is further emphasized by the directionality of the gray arrows, which indicate direction of the net flux between each forward/reverse enzyme pair. These findings suggest an early, rapid dismantling of certain “higher□order” sphingolipid pools, presumably to fuel signaling processes mediated by free bases.

**Figure 6.**
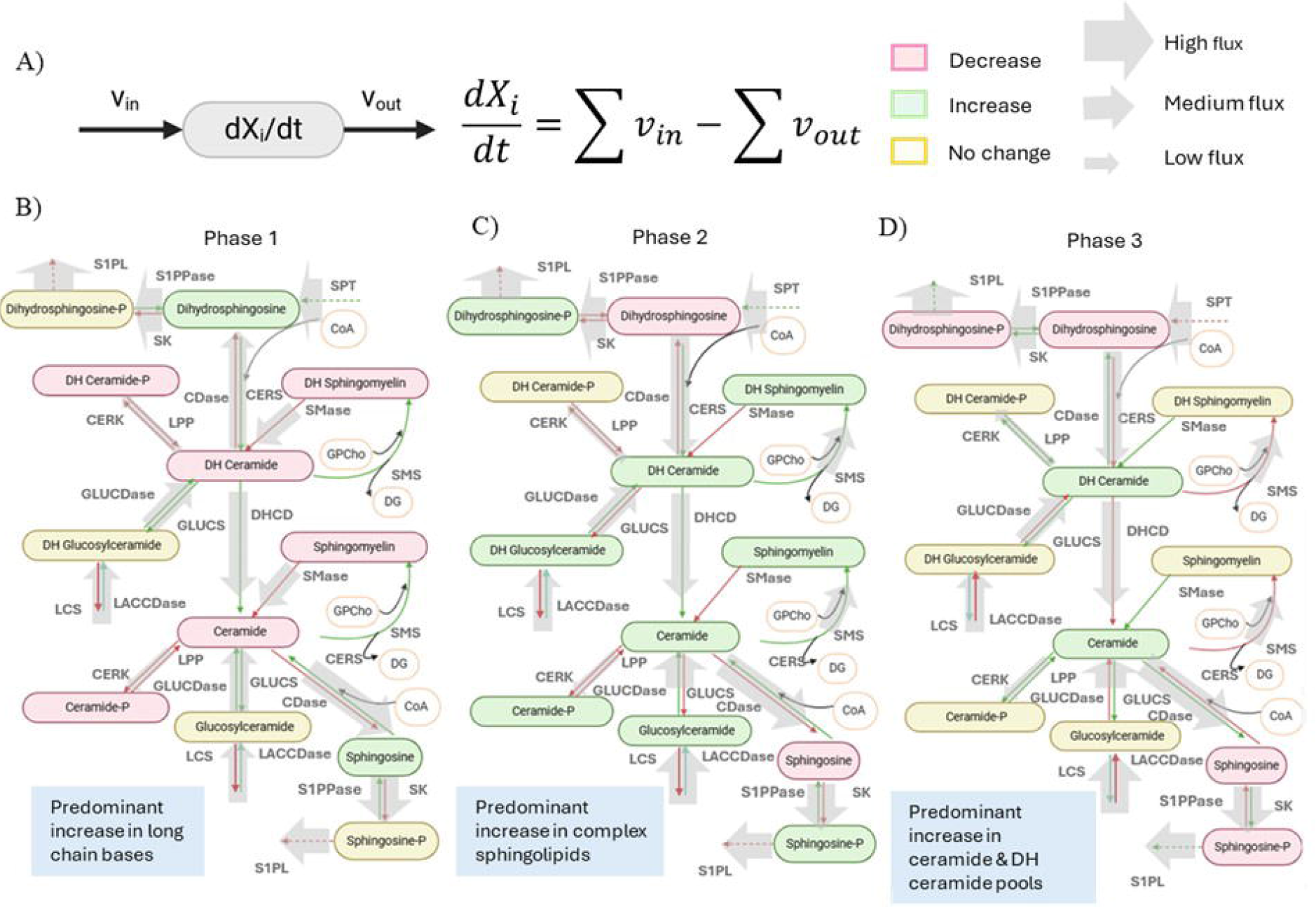
Coordinated flux changes drive shifts in sphingolipid concentrations over three phases of KdO□□Lipid A stimulation. **(A–C)** Network diagrams for **(A)** Phase□1, **(B)** Phase□2, and **(C)** Phase□3, depicting how lipid concentrations and fluxes change over time. Lipid boxes are color□coded as follows: green for increased concentration, red for decreased, and yellow for unchanged. Arrows are likewise colored to indicate increased (green), decreased (red), or unchanged (yellow) flux. Gray arrows denote net reaction directions for each reaction pair, with arrow thickness proportional to flux magnitude. In Phase□1, mass predominantly flows from complex sphingolipids (e.g., sphingomyelin, ceramide 1 phosphate, ceramide) toward long□chain bases (sphingosine, dihydrosphingosine). In Phase□2, complex sphingolipid pools are rebuilt and expanded, while accumulated long□chain bases are converted into their phosphorylated forms. Finally, in Phase□3, remaining long□chain bases and base phosphates are removed, concurrent with continued expansion of the ceramide pool.

By Phase 2 (4–12 h), the metabolic emphasis shifts. Many complex sphingolipids (e.g., sphingomyelin, glucosylceramide) rebound or expand (**Figure 6C**, green boxes), supported by increased flux through synthases (e.g., SMS, GLUCERS). Meanwhile, a portion of the accumulated long□chain bases is channeled into their phosphorylated forms (SphP, DHSphP). These phosphorylated bases also appear in green for flux or concentration in certain nodes, reflecting ongoing pro□survival or pro□migratory signaling. The data suggest that macrophages actively restore higher□order lipid pools while diversifying base intermediates via phosphorylation—possibly to balance pro□inflammatory and reparative processes.

In Phase 3 (12–24 h), most long□chain bases and their phosphorylated derivatives are downregulated (red boxes/arrows in **Figure 6D**), implying a partial resolution of the early and intermediate accumulations. Many of the complex sphingolipids (e.g. sphingomyelin and glucosylceramide) stabilize at new elevated levels (yellow boxes) Notably, ceramide and dihydroceramide remain elevated or continue to expand (green), suggesting that these pools either are still actively synthesized or have not yet returned to baseline. This ongoing ceramide enrichment could underlie persistent pro□inflammatory signaling—or a preparation phase for further membrane remodeling—in late□stage inflammation. Overall, the phase□specific patterns provide a system□level view of how core sphingolipid pools ebb and flow through each stage of the macrophage response. Such coordinated flux modulations help explain how macrophages finely tune sphingolipid levels for diverse functional outcomes—ranging from cell signaling to membrane biogenesis—over the full inflammatory timeline.

### Key Enzymatic Fluxes Drive Major Shifts in Sphingolipid Concentrations

The complexity of sphingolipid metabolism results in multiple control strategies for the sphingolipid concentrations. For ceramide, which showed persistent elevation through Phase 3, control of concentration seems to be distributed across multiple fluxes: First, sphingomyelin synthase (SMS) and glucosylceramidase (GLUCDase) show by far the largest changes in flux in the stimulated group compared to the control group (**Figure 7A,B**). Because GLUCDase produces ceramide and SMS consumes ceramide, the net effect is the flow of mass from GLUCER to SM. These two large fluxes usually balance each other out, but SMS dominates during Phase 2 resulting in a net loss of ceramide. At the same time, the other smaller fluxes such as dihydroceramide desaturase (DHCD) contribute mass to the ceramide pool resulting in a net increase in concentration. This suggests that ceramide accumulation during inflammation stems from a balance of the loss of mass during flow from glucosylceramide to sphingomyelin and the gain of mass from synthetic pathways. Notably, these results suggest that a small change in SMS or GluCDase would disrupt the balance between these two reactions and allow for significant changes in ceramide dynamics.

**Figure□7.**
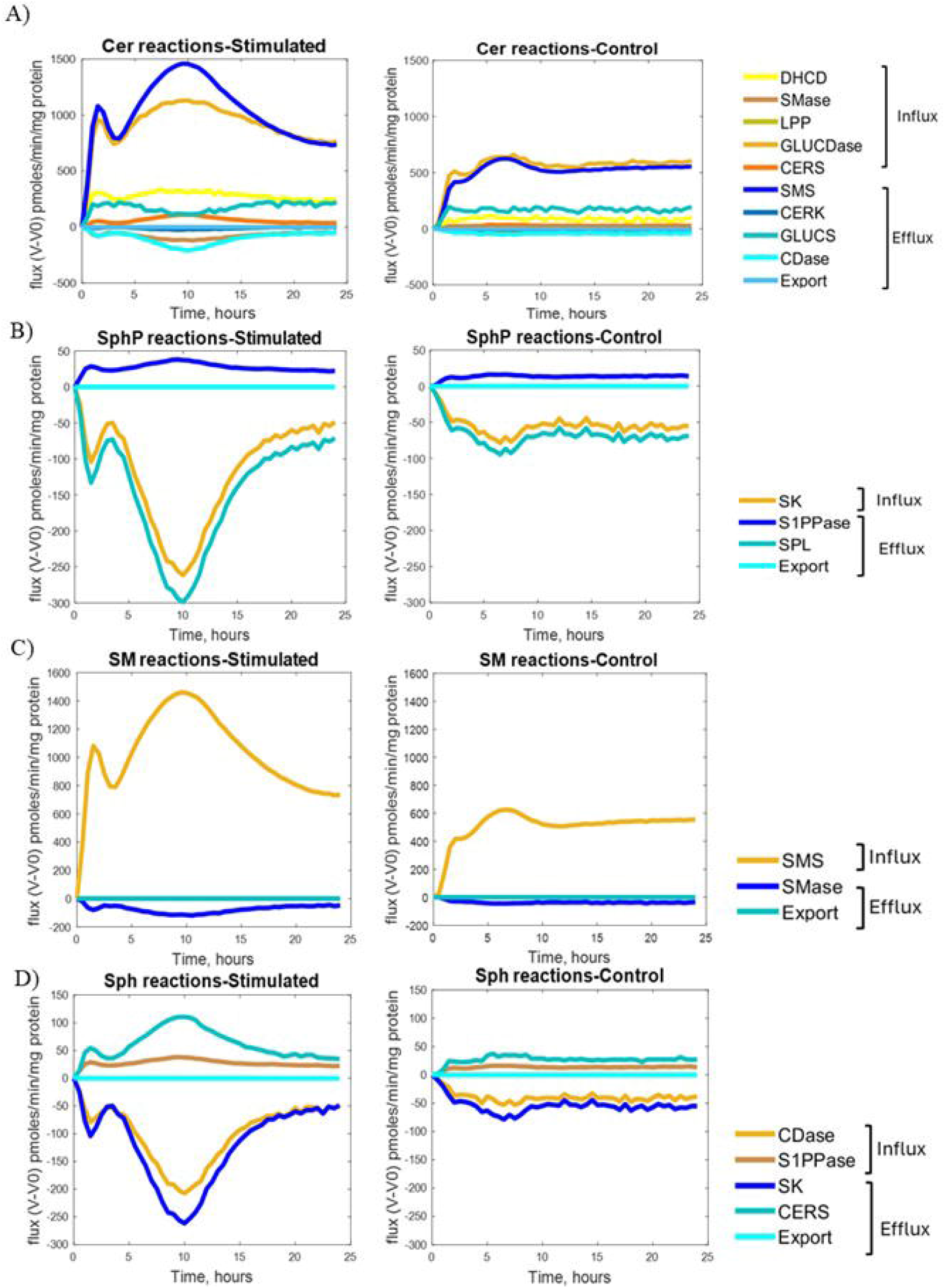
A few “key” fluxes dominate sphingolipid concentration changes after KdO□□Lipid□A stimulation. **(A)** Ceramide (Cer) levels are primarily regulated by sphingomyelin synthase (SMS) and glucosylceramidase (GLUCDase) fluxes. **(B)** Sphingosine phosphate (SphP) concentrations hinge on sphingosine kinase (SK) and sphingosine□1□phosphate lyase (SPL) activities. **(C)** Sphingomyelin (SM) concentrations are largely determined by changes in SMS flux. **(D)** Sphingosine (Sph) concentrations respond to SK and ceramidase (CDase) fluxes. In each panel, flux trajectories are shown as deviations from their baseline (t□=□0) values. Notably, despite the broader complexity of sphingolipid metabolism, a small subset of enzymatic reactions appears to drive most of the observed concentration shifts in these four metabolites. This finding underscores the potential for targeted modulation of just a few fluxes—rather than tackling the entire pathway—to achieve substantial changes in sphingolipid homeostasis and macrophage inflammatory response.

Sphingosine-1-phosphate (SphP) levels, critical for pro-survival signaling, are tightly controlled by the opposing actions of just 2 enzymes: sphingosine kinase (SK) and sphingosine-1-phosphate lyase (SPL) (**Figure 7B**). The substantial drop in SPL flux during Phase 2 allows for significant accumulation of SphP, potentially moderating pro-survival signals as the inflammatory response progresses. This finding reveals a previously unappreciated temporal control point in sphingolipid-mediated inflammation.

Sphingomyelin concentrations track almost exclusively with SMS flux (**Figure 7C**), highlighting this enzyme as a master regulator of membrane sphingolipid composition during inflammation. The dramatic increase in SMS flux during Phase 2 suggests extensive membrane remodeling coincides with the intermediate inflammatory response. Meanwhile, sphingosine levels are primarily regulated by the coordinated activities of SK and ceramidase (**Figure 7D**), with their fluxes showing strong temporal correlation but opposite directions during Phase 2. Similar patterns of concentrations being dominated by a few key fluxes emerged for other sphingolipids as well (**Figure S4**). The dominant fluxes for all sphingolipids are summarized in **Table 6**.

**Table 6.** Dominant Fluxes for Each Sphingolipid

| No. | Sphingolipid | Dominant Fluxes |
| --- | --- | --- |
| 1. | Dihydrosphingosine | SPT, CDase |
| 2. | Dihydrosphingosine Phosphate | SPL, SK |
| 3. | Dihydroceramide | GLUCDase, SMS |
| 4. | Dihydrosphingomyelin | SMS |
| 5. | Dihydroceramide Phosphate | CERK, LPPase |
| 6. | Dihydroglucosylceramide | GLUCDase |
| 7. | Ceramide | GLUCDase, SMS |
| 8. | Sphingomyelin | SMS |
| 9. | Ceramide Phosphate | CERK, LPP |
| 10. | Glucosylceramide | LACCDase, GLUCDase |
| 11. | Sphingosine | SK |
| 12. | Sphingosine Phosphate | SPL |

These findings reveal that while the sphingolipid network contains numerous interconnected reactions, therapeutic targeting of just a few key enzymes - particularly SMS, GluCDase, SK, or SPL - might be sufficient to reshape the inflammatory sphingolipid landscape. The temporal profiles of these dominant fluxes also suggest that the timing of such interventions could be crucial, with Phase 2 representing a particularly dynamic window for metabolic modulation.

### Sensitivity Analysis Identifies Critical Regulatory Points in the Sphingolipid Network

To understand how perturbations in enzyme activities influence sphingolipid levels, we performed dynamic sensitivity analysis. Based on this analysis, we produced sensitivity matrices mapping the sensitivities of lipid concentrations to changes in enzyme activities at 0, 4, and 12 hours post KLA stimulation. (**Figure 8A-C**). The matrices reveal that at a given time point each lipid responds strongly to only a few of the enzymes in the network. However, those influential lipids change as time progresses. This pattern suggests that targeting specific enzymes in specific time windows could achieve selective control over particular sphingolipid species. Ceramide sensitivity analysis illustrates this principle (**Figure 8B**). Among the network’s enzymes, LacCDase shows the highest sensitivity coefficient for ceramide at all time points, peaking at 10 hours, while other enzymes like CDase and DHCD show modest sensitivities only in Phase 1and SMS shows modest sensitivity only in Phase 2. This indicates that ceramide levels could be effectively modulated through LacCDase activity. This is consistent with the results from Figure 7, which identified GLUCDase as a key flux into the ceramide pool because changes in flux from LACCDase must flow through GLUCDase to reach ceramide. To test this prediction, we simulated progressive inhibition of LacCDase at 10 hours, the time identified to have the highest sensitivity (**Figure S5**). The model shows that 24 hour ceramide concentrations decrease steadily with increasing LacCDase inhibition at 10 hours, although the concentration is never able to completely reach the control level. This finding suggests that partial enzyme inhibition might be sufficient to reduce elevated ceramide levels observed during inflammation. The sensitivity matrix also reveals unexpected network relationships. For instance, dihydrosphingosine-based species show distinct sensitivity patterns compared to their sphingosine-based counterparts, suggesting separate regulatory mechanisms for these precursor pools. These insights from our computational analysis provide a quantitative framework for predicting how targeted enzymatic interventions might propagate through the sphingolipid network.

**Figure□8.**
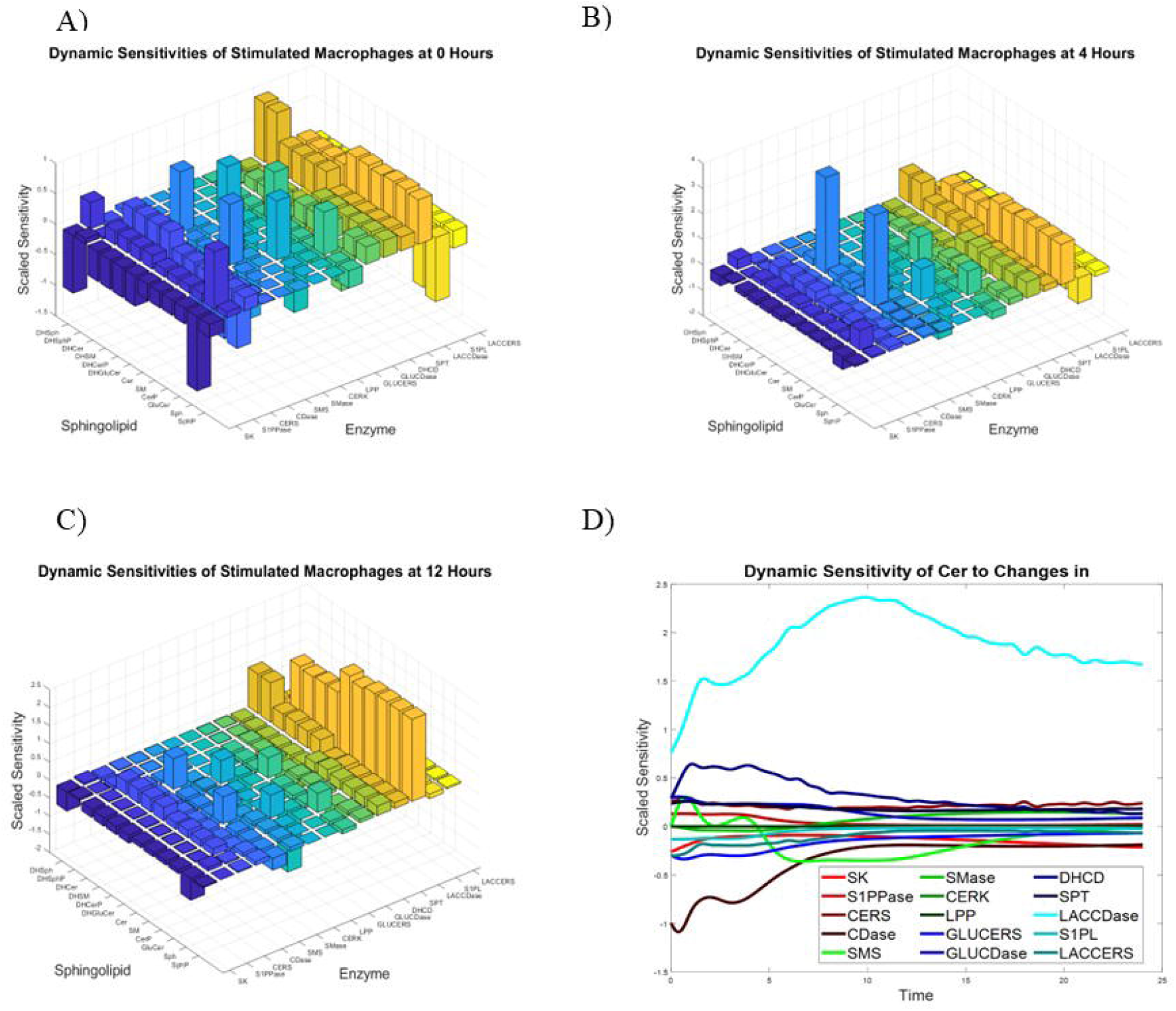
Sensitivities of Sphingolipids to Enzymes Change over Time, but a Small Number of Enzymes Dominate. The scaled sensitivities of sphingolipid concentrations with respect to enzyme activities were calculated at **(A)** 0 hours, **(B)** 4 hours, and **(C)** 12 hours post KLA stimulation. **(D)** Sensitivity of ceramide concentration to changes in enzyme activities throughout the entire 24-hour experiment post KLA stimulation.

### Integration of Sphingolipid Dynamics with Macrophage Functional States

Our temporal analysis of sphingolipid metabolism reveals distinct metabolic programs that align with key functional transitions in macrophage activation (**Figure 9**). In Phase 1 (0-4h), the selective increase in sphingosine and dihydrosphingosine, known inhibitors of protein kinase C (PKC), suggests an initial regulatory checkpoint that temporarily constrains full inflammatory activation[1]. This early phase shows minimal changes in other sphingolipid species, indicating that cells maintain near-baseline membrane composition while establishing appropriate activation thresholds. Phase 2 (4-12h) marks a dramatic metabolic shift characterized by enhanced membrane production and vesicular trafficking. The coordinated upregulation of complex sphingolipids, including ceramide, dihydroceramide, and glucosylceramide, supports increased endolysosomal activity[6].

**Figure 9.**
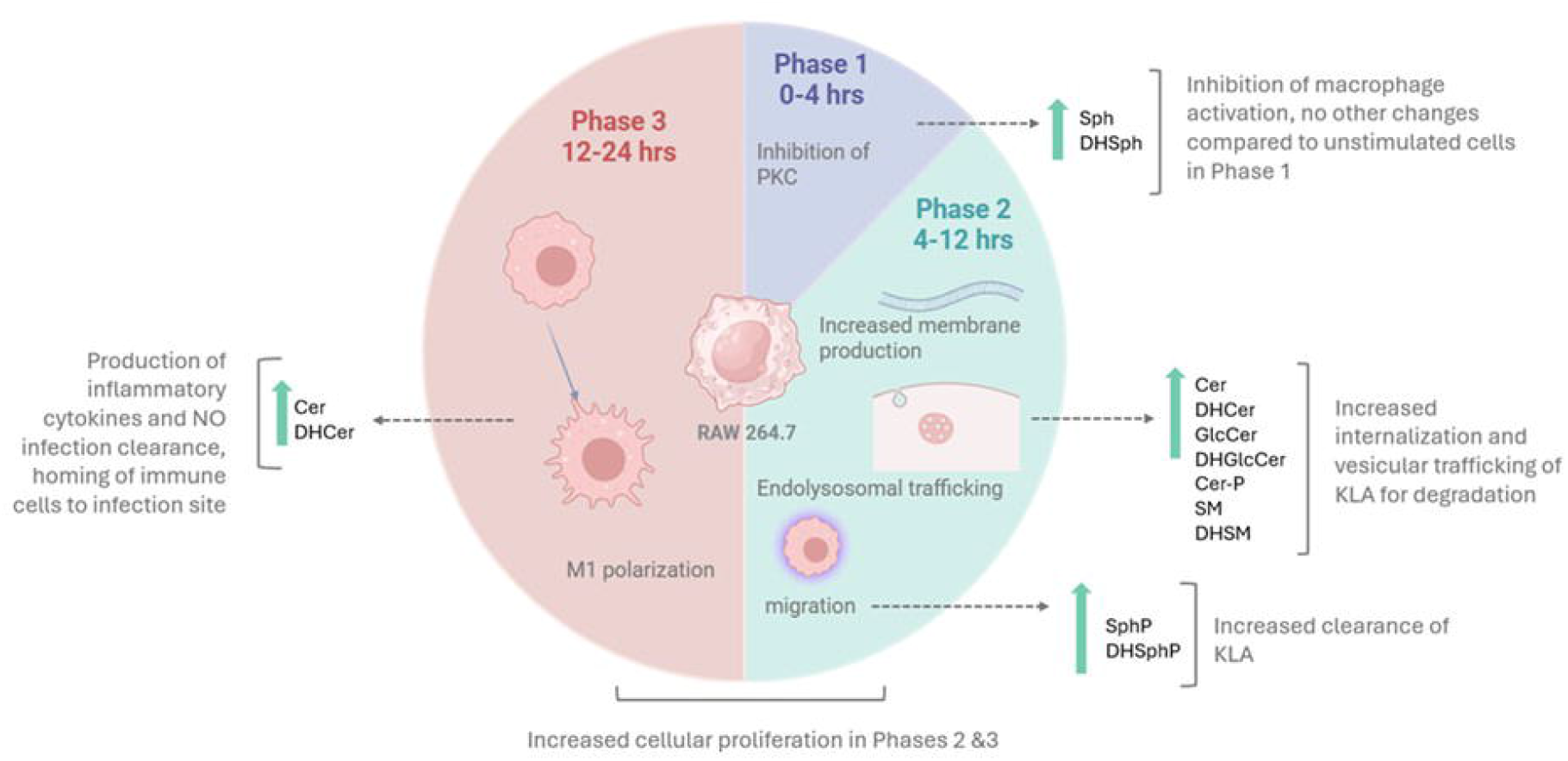
Three□phase macrophage response to KdO□□Lipid□A involves dynamic lipid remodeling and functional adaptations. Phase□1 (0–4□h) features relatively limited changes in macrophage activation alongside the inhibition of PKC, with cells remaining close to baseline in both morphology and sphingolipid levels. Phase□2 (4–12□h) is characterized by increased membrane production and pronounced endolysosomal trafficking, accompanied by upregulated sphingolipid species (e.g., Cer, DHCer, GlcCer, and others). This phase supports enhanced internalization and vesicular processing of KLA for downstream immune signaling. Phase□3 (12–24□h) culminates in robust M1 polarization, with elevated Cer and DHCer contributing to the production of inflammatory cytokines, nitric oxide (NO), and the recruitment of additional immune cells. Meta□analysis of the literature and our data indicates that each phase integrates distinct metabolic programs to drive effective responses against inflammatory stimuli. The sequential upregulation of sphingolipids during Phases□2 and□3 supports increased cellular proliferation, membrane remodeling, and ultimately, inflammatory effector functions essential for infection clearance.

This metabolic remodeling also coincides with elevated S1P and dhS1P levels, which promote cell survival and migration[13]. Meta-analysis of the LIPID MAPS dataset reveals increased DNA content during this phase, suggesting that sphingolipid changes also support heightened cellular proliferation[20]. By Phase 3 (12-24h), the sphingolipid landscape shifts toward an inflammatory profile dominated by ceramide and dihydroceramide accumulation. This metabolic signature aligns with M1-like polarization[28], enabling enhanced production of inflammatory mediators and recruitment of additional immune cells. The sustained elevation of these pro-inflammatory sphingolipid species, even as other metabolites begin returning to baseline, suggests their importance in maintaining appropriate immune activation[15].This three-phase model demonstrates how macrophages orchestrate sequential sphingolipid programs to achieve effective inflammatory responses while maintaining essential cellular functions. The temporal coordination between lipid remodeling and functional adaptations provides new insight into the metabolic regulation of innate immunity[6].

## Discussion

Our systematic analysis of sphingolipid metabolism in activated macrophages reveals a coordinated three-phase response that provides new mechanistic insights into inflammatory regulation. The temporal evolution from early sphingolipid mobilization to later partial resolution demonstrates how macrophages orchestrate metabolic programs to support distinct functional states.

Phase 1 (0–4 h) is marked by an increase in sphingosine and dihydrosphingosine, two long□chain bases that can inhibit protein kinase C and thus transiently dampen TLR4□mediated signaling [1, 11, 29]. This early checkpoint could serve as a regulatory buffer before full-scale macrophage activation. By Phase 2 (4–12 h), we observe pronounced accumulations of complex sphingolipids (e.g., sphingomyelin, glucosylceramide) and the bioactive lipids S1P and DHS1P, potentially linked to membrane biogenesis and endolysosomal trafficking [20, 30]. Enhanced S1P levels in particular are associated with cell survival, migratory signaling, and anti□apoptotic pathways [15, 31]. Finally, Phase 3 (12–24 h) is typified by ceramide and dihydroceramide accumulation, reminiscent of an M1□like pro□inflammatory phenotype [28]. Although many metabolites and fluxes partially return toward baseline, ceramide elevations often persist, suggesting incomplete resolution that could maintain a heightened inflammatory tone if not further regulated [7, 11].

One emerging concept is that macrophages can retain a “metabolic memory” of prior inflammatory signals, influencing how they respond to subsequent challenges [6]. Our data illustrate that certain sphingolipid pools (e.g., ceramide) and enzymes remain altered into Phase 3, possibly priming macrophages for heightened responses if stimulation recurs. Intriguingly, anti□inflammatory cytokines such as IL□10 have been shown to constrain sphingolipid metabolism and reduce excessive inflammatory signaling [29, 30]. It is plausible that IL□10 or related signals might tip the balance of late□phase sphingolipid metabolism back toward S1P or ceramide□1□phosphate, fostering a more resolving or reparative macrophage state [8, 31]. Future experiments manipulating IL□10 levels or downstream effectors in conjunction with time□resolved sphingolipid profiling could clarify how these regulatory axes influence “metabolic memory” within the macrophage.

Our sensitivity analysis suggests that targeting key enzymes (e.g., lactosylceramidase, sphingomyelin synthase) at different phases can exert outsized impacts on sphingolipid concentrations. This aligns with recent findings that sphingolipid metabolism is a druggable pathway for modulating inflammation [16, 32]. Importantly, the timing of such interventions appears crucial. In Phase 2, for example, the surge in S1P and other phosphorylated bases may present an optimal window to either attenuate pro□survival signals or enhance resolving mechanisms—depending on the clinical context [33, 34]. Conversely, inhibiting ceramide synthesis or conversion in Phase 3 could help prevent chronic inflammation and tissue damage. By providing a quantitative and temporal perspective, our approach opens the door to designing phase□aligned therapies where enzyme inhibition or activation is timed to the metabolic transitions most likely to influence outcome. Similar methods were used by Alvarez-Vasquez to create a biochemical systems theoretical model of sphingolipid metabolism in Saccharomyces cerevisiae to predict the effects of flux modulations, genetic enzyme perturbations and the impact of inositol regulation with high agreement to experimental results. Importantly, the model also allowed the simulation of changes in fatty-acid precursors that are difficult or test experimentally, thereby demonstrating its utility in in-silico “thought experiments”[35]. Recent studies have highlighted the complexity of immunometabolism, with macrophages integrating glycolysis, fatty acid oxidation, arginine metabolism, and sphingolipid pathways to fine□tune pro□ vs. anti□inflammatory phenotypes [1, 15]. Our observation that S1P and ceramide fluxes strongly shape cell fate decisions underscores the synergy between lipid remodeling and classical immune signaling. Moreover, the partial resolution we observed after 24 hours may reflect macrophages striving for a balanced inflammatory response—strong enough to neutralize pathogens yet not so excessive as to induce bystander tissue damage [6, 31]. Understanding how these sphingolipid programs intersect with other metabolic circuits, including the TCA cycle or pentose phosphate pathway, represents a significant next step [16].

## Limitations

Despite offering new insights, our study has certain limitations. First, enzyme activities were inferred from literature sources rather than measured directly in RAW 264.7 macrophages; thus, specific reaction rates may differ in vivo or under varied culture conditions [10]. Second, our bulk cell analysis cannot capture potential subcellular compartmentalization of sphingolipid metabolism or cell-to-cell heterogeneity. Recent advances in single-cell lipidomics using trapped ion mobility separation [36] and spatial mapping techniques [14, 21] provide valuable insights into the spatial distribution and heterogeneity of sphingolipid dynamics during macrophage activation, and explore how different macrophage subpopulations might exhibit distinct metabolic programs during inflammation [37]. Third, macrophages in tissues encounter multicellular contexts, gradients of cytokines, and dynamic oxygen levels—all absent in simplified in vitro systems. Future work could examine how local sphingolipid gradients (e.g., in fibrotic lung, atherosclerotic plaques or traumatic tissue loss) shape macrophage responses in vivo [3]. Fourth, although we explored a range of flux behaviors, additional multi□omics analyses (e.g., transcriptomics, proteomics) might further elucidate how sphingolipid metabolism integrates with other immunometabolic processes. Nonetheless, our model provides a valuable starting point for predictive analyses, bridging in vitro measurements with potential in vivo applications [16, 34].

## Conclusions

Together, these findings establish a quantitative foundation for understanding how macrophages coordinate sphingolipid metabolism during inflammation. By delineating a three□phase progression—early dampening (Phase 1), complex sphingolipid/ S1P buildup (Phase 2), and partial resolution with persistent ceramide (Phase 3)—we highlight temporal niches in which therapeutic interventions could be most effective. The identification of phase□specific control points (e.g., glucosylceramidase, sphingosine kinase) and metabolite “memory” (e.g., sustained ceramide elevations) has direct implications for designing more targeted anti□inflammatory therapies. Moving forward, integrating these insights with complementary immunometabolic data and in vivo validation may yield robust strategies to modulate macrophage□driven inflammation without broadly inhibiting their essential immune functions.

## List of Sphingolipid Abbreviations

Sph: Sphingosine
Sph P: Sphingosine 1 Phosphate
DHSph: Dihydrosphingosine
Sph P: Dihydrosphingosine 1 Phosphate
Cer: Ceramide
DHCer: Dihydroceramide
Cer P: Ceramide 1 Phosphate
DHCer P: Dihydroceramide 1 Phosphate
GluCer: Glucosylceramide
DHGluCer: Dihydroglucosylceramide
SM: Sphingomyelin
DHSM: Dihydrosphingomyelin
LacCer: Lactosylceramide

