## Supplemental Figures for "Resolving vs. Non-resolving Sphingolipid Dynamics During Macrophage Activation: A Time-resolved Metabolic Analysis"

### Slide 1
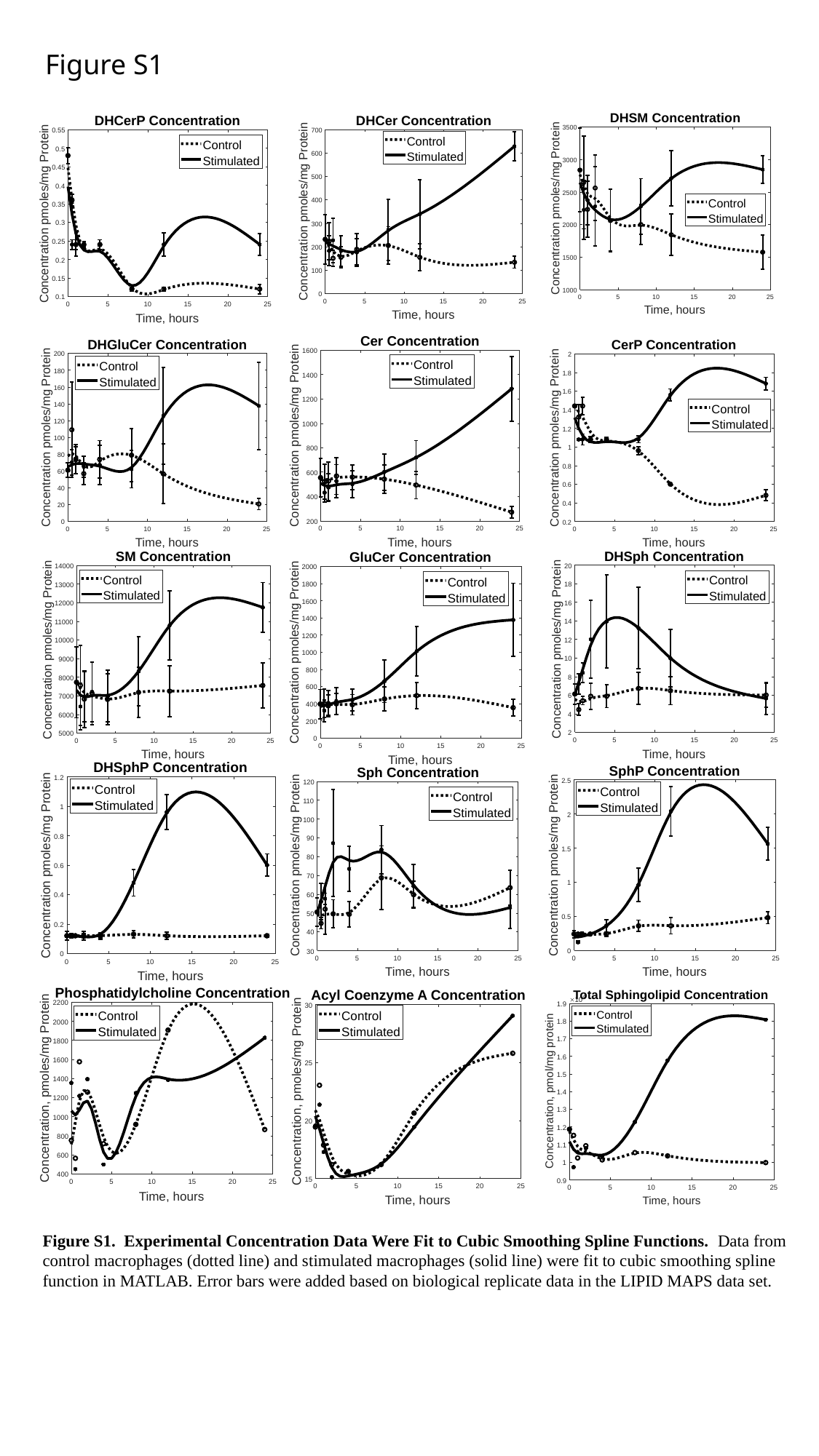

Figure S1
Figure S1. Experimental Concentration Data Were Fit to Cubic Smoothing Spline Functions. Data from control macrophages (dotted line) and stimulated macrophages (solid line) were fit to cubic smoothing spline function in MATLAB. Error bars were added based on biological replicate data in the LIPID MAPS data set.

### Slide 2
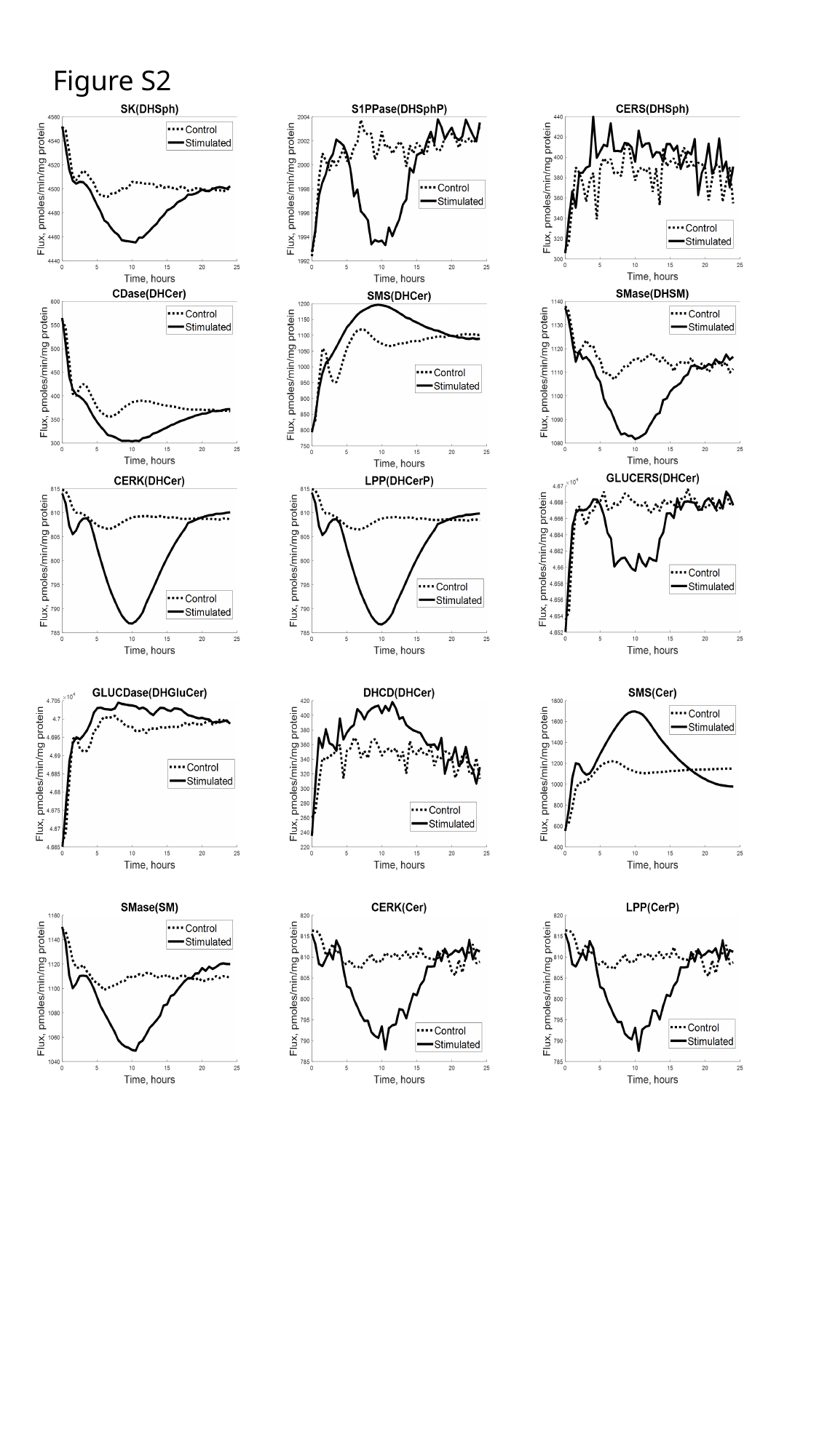

Figure S2

### Slide 3
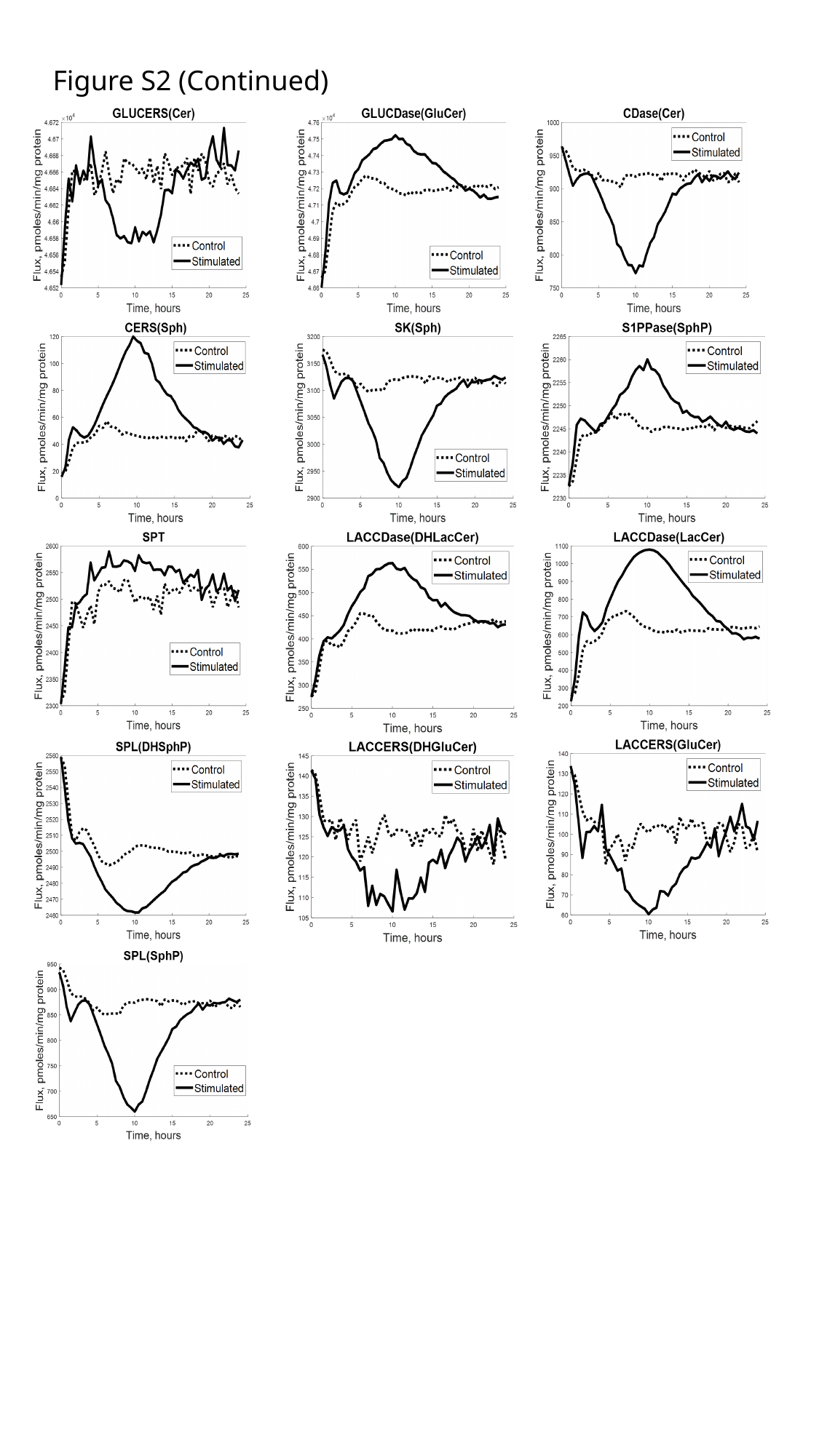

Figure S2 (Continued)

### Slide 4
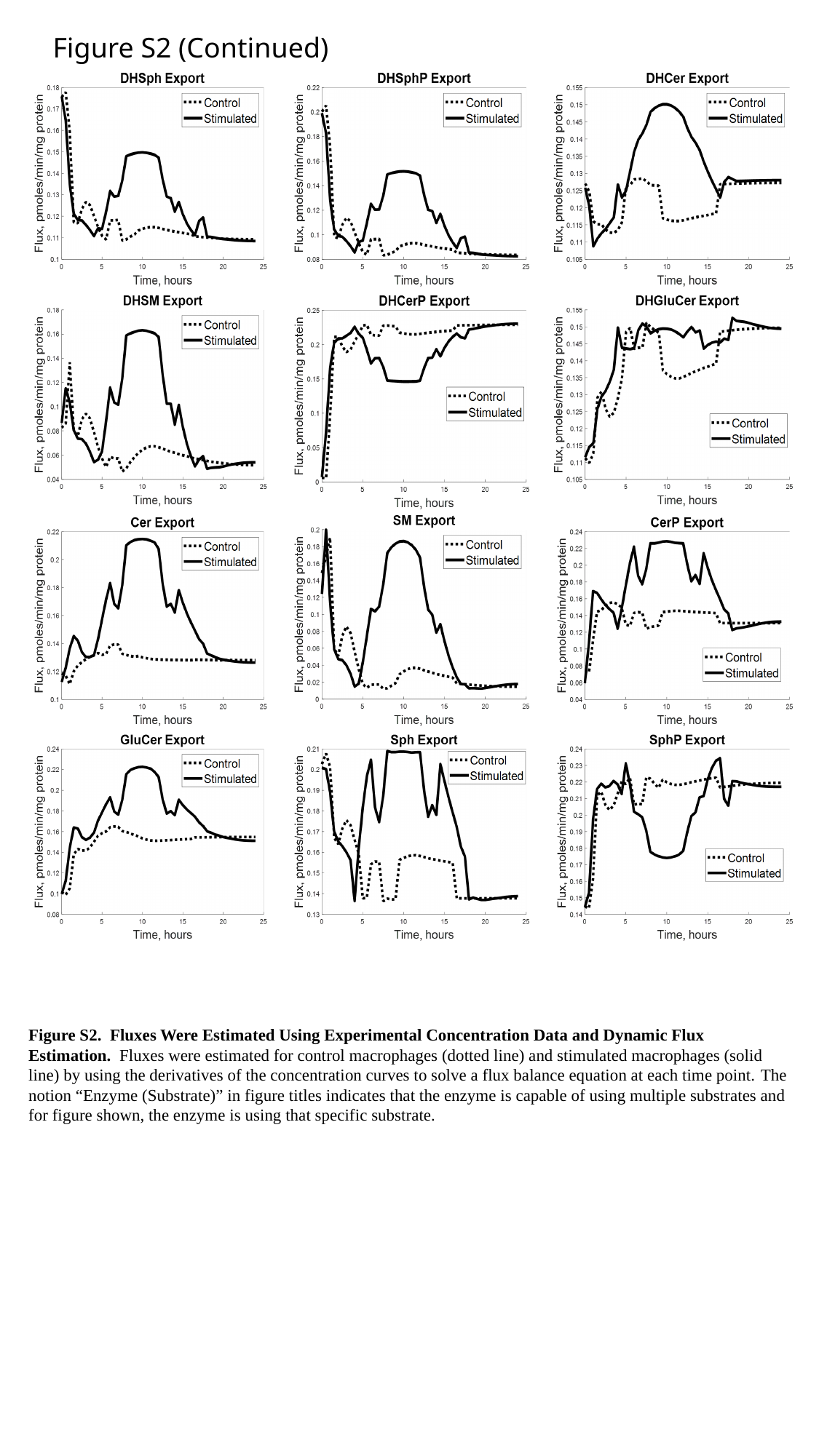

Figure S2 (Continued)
Figure S2. Fluxes Were Estimated Using Experimental Concentration Data and Dynamic Flux Estimation. Fluxes were estimated for control macrophages (dotted line) and stimulated macrophages (solid line) by using the derivatives of the concentration curves to solve a flux balance equation at each time point. The notion “Enzyme (Substrate)” in figure titles indicates that the enzyme is capable of using multiple substrates and for figure shown, the enzyme is using that specific substrate.

### Slide 5
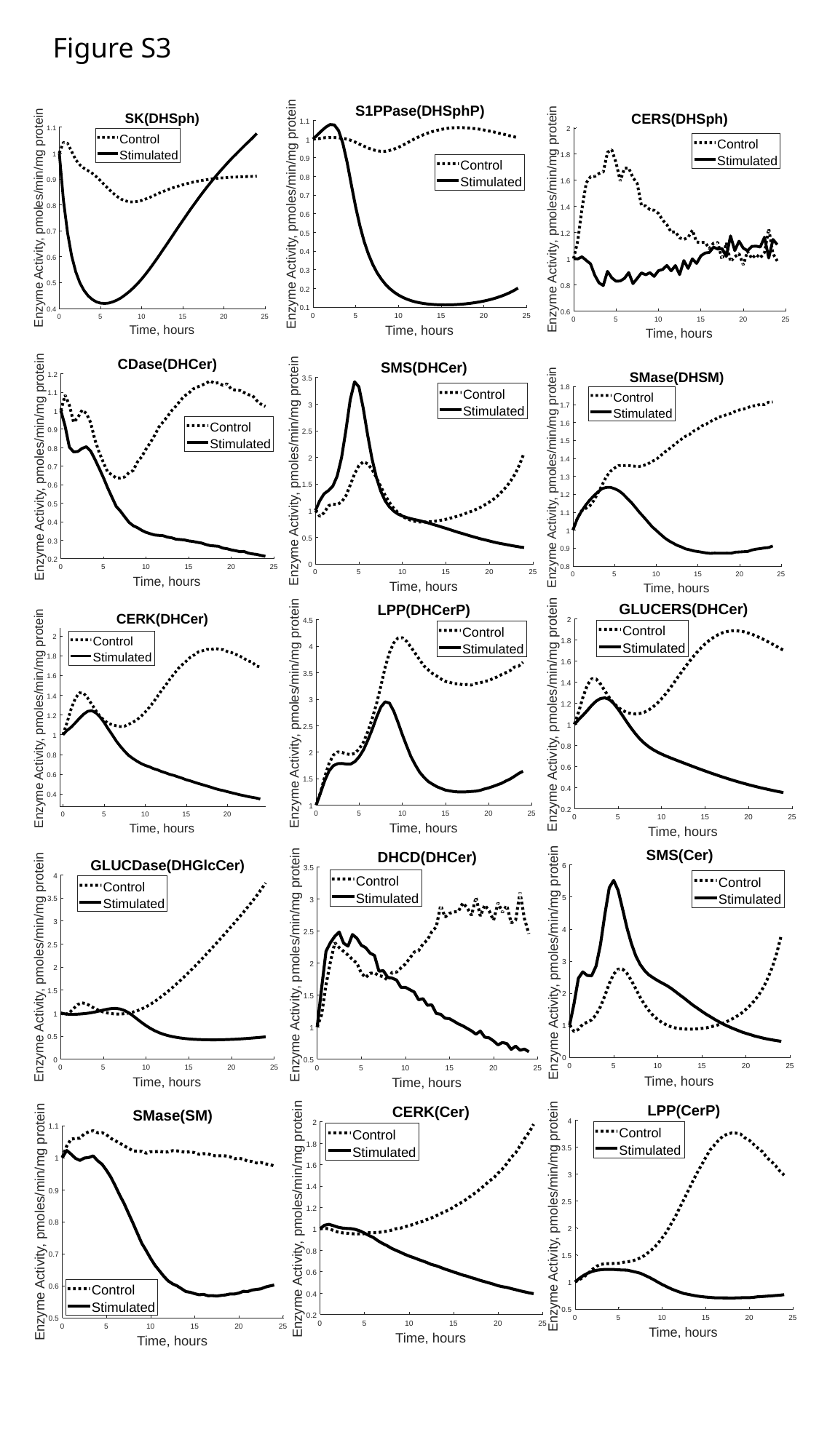

Figure S3

### Slide 6
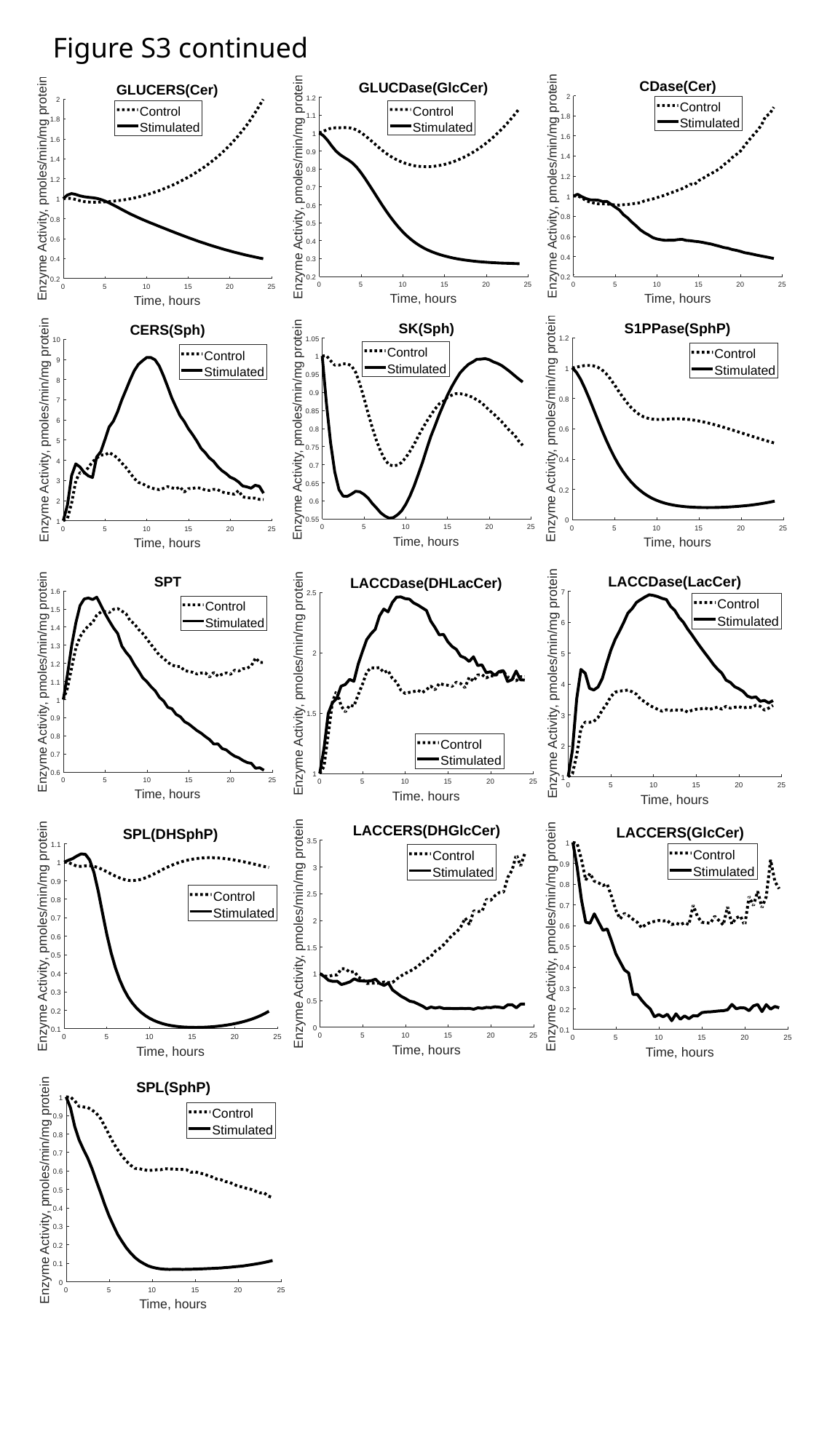

Figure S3 continued

### Slide 7
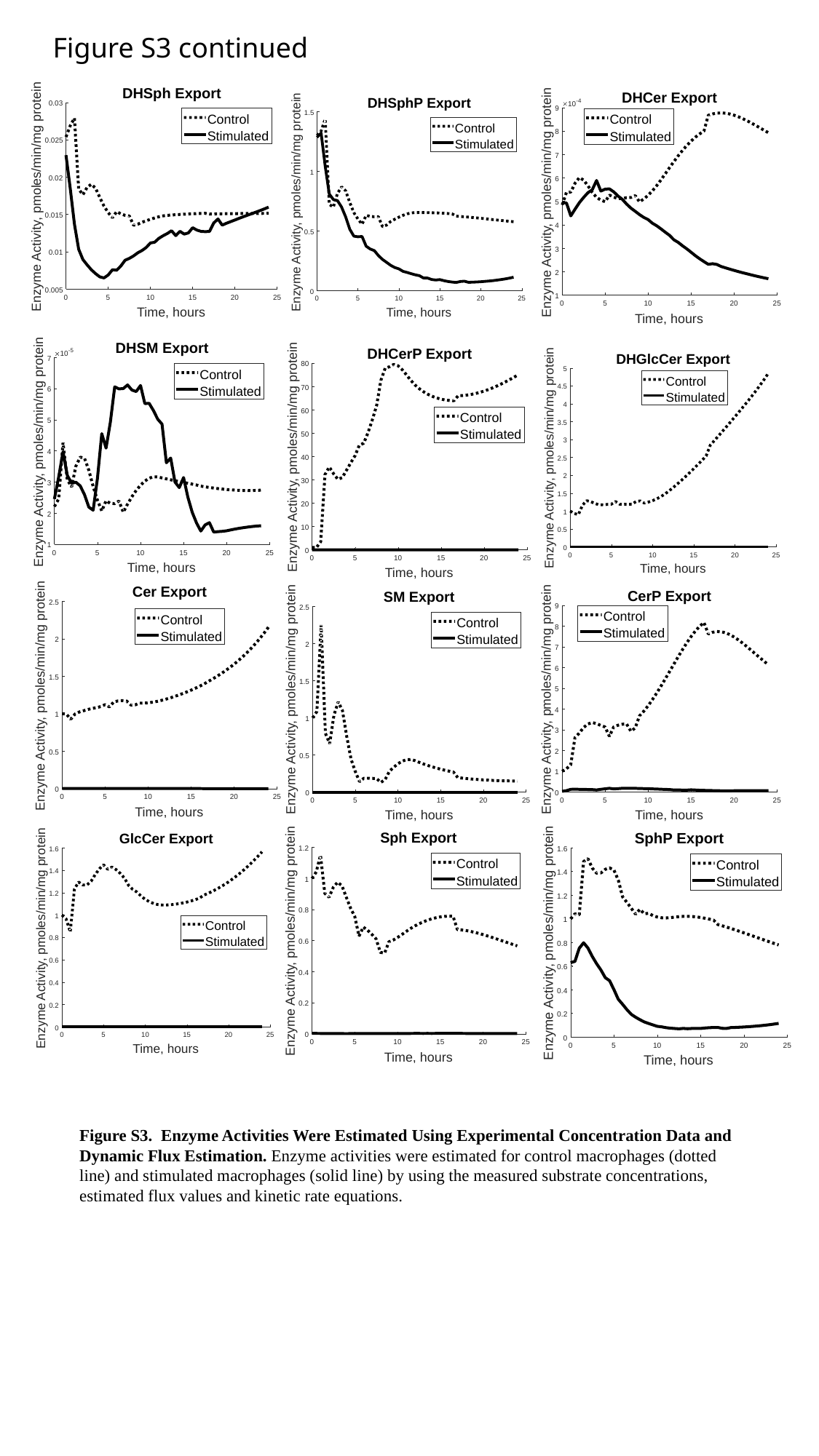

Figure S3 continued
Figure S3. Enzyme Activities Were Estimated Using Experimental Concentration Data and Dynamic Flux Estimation. Enzyme activities were estimated for control macrophages (dotted line) and stimulated macrophages (solid line) by using the measured substrate concentrations, estimated flux values and kinetic rate equations.

### Slide 8
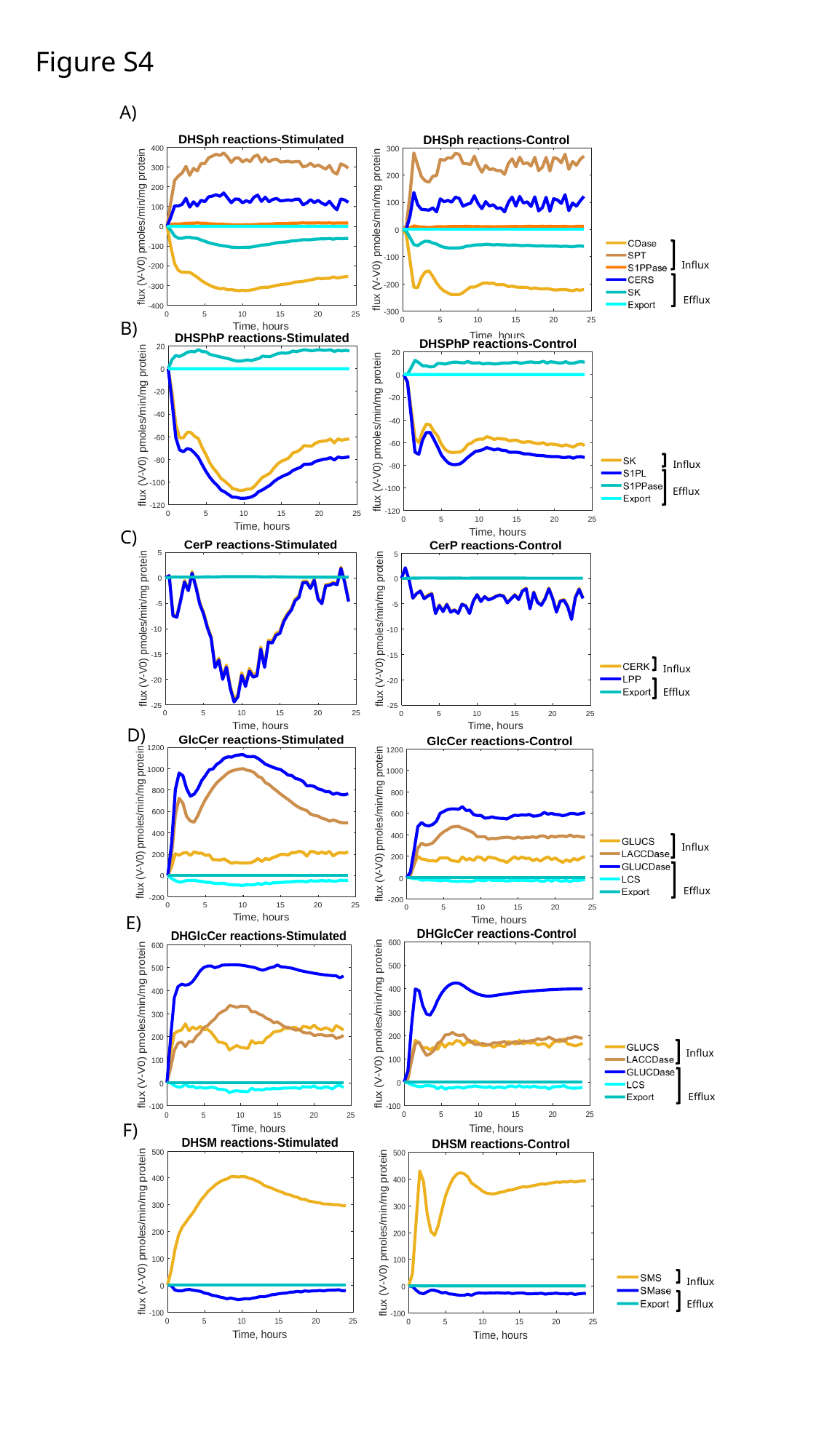

Figure S4
A)
Influx
Efflux
B)
Influx
Efflux
C)
Influx
Efflux
D)
Influx
Efflux
E)
Influx
Efflux
F)
Influx
Efflux

### Slide 9
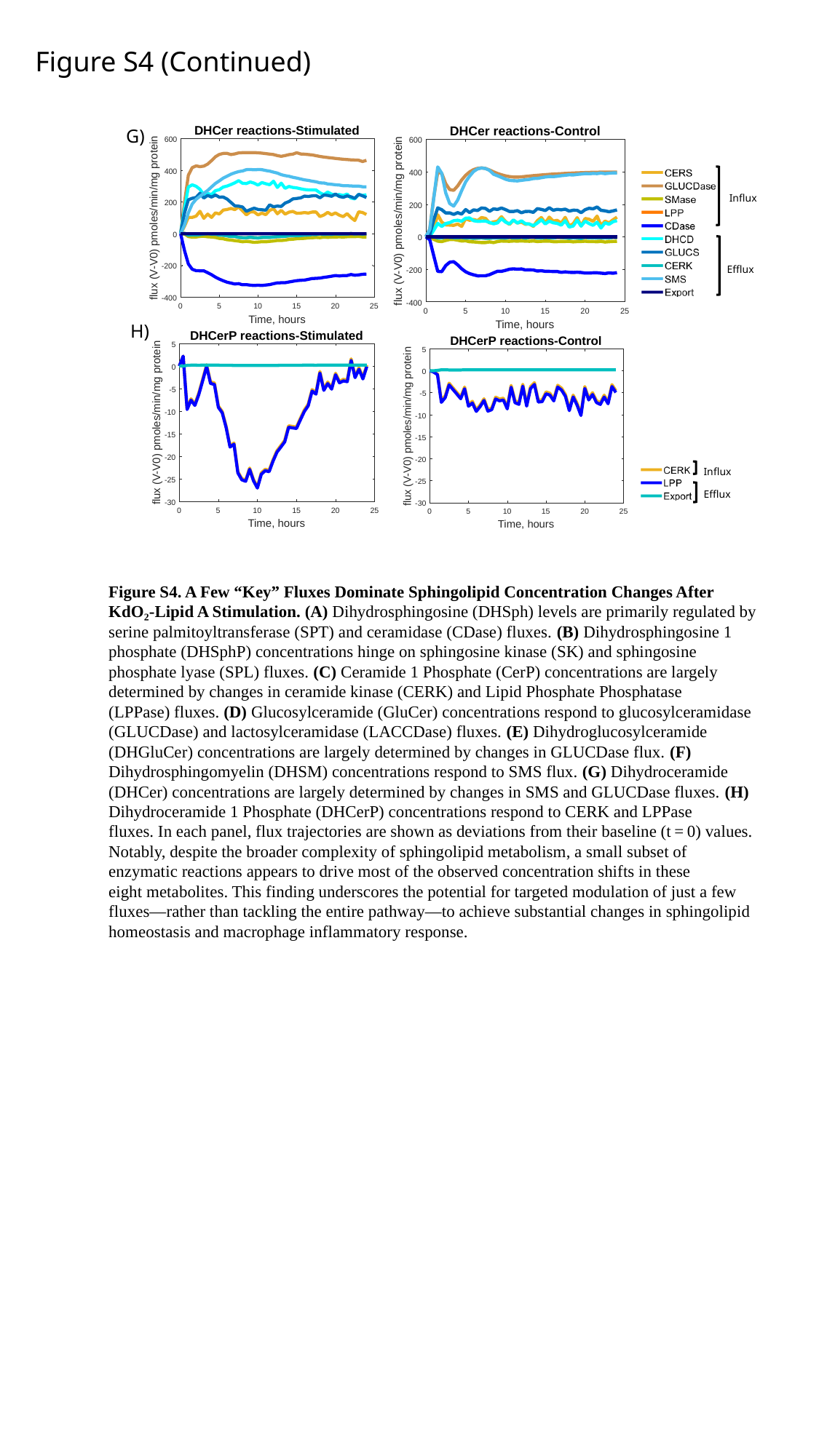

Figure S4 (Continued)
G)
Influx
Efflux
H)
Influx
Efflux
Figure S4. A Few “Key” Fluxes Dominate Sphingolipid Concentration Changes After KdO₂‐Lipid A Stimulation. (A) Dihydrosphingosine (DHSph) levels are primarily regulated by serine palmitoyltransferase (SPT) and ceramidase (CDase) fluxes. (B) Dihydrosphingosine 1 phosphate (DHSphP) concentrations hinge on sphingosine kinase (SK) and sphingosine phosphate lyase (SPL) fluxes. (C) Ceramide 1 Phosphate (CerP) concentrations are largely determined by changes in ceramide kinase (CERK) and Lipid Phosphate Phosphatase (LPPase) fluxes. (D) Glucosylceramide (GluCer) concentrations respond to glucosylceramidase (GLUCDase) and lactosylceramidase (LACCDase) fluxes. (E) Dihydroglucosylceramide (DHGluCer) concentrations are largely determined by changes in GLUCDase flux. (F) Dihydrosphingomyelin (DHSM) concentrations respond to SMS flux. (G) Dihydroceramide (DHCer) concentrations are largely determined by changes in SMS and GLUCDase fluxes. (H) Dihydroceramide 1 Phosphate (DHCerP) concentrations respond to CERK and LPPase fluxes. In each panel, flux trajectories are shown as deviations from their baseline (t = 0) values. Notably, despite the broader complexity of sphingolipid metabolism, a small subset of enzymatic reactions appears to drive most of the observed concentration shifts in these eight metabolites. This finding underscores the potential for targeted modulation of just a few fluxes—rather than tackling the entire pathway—to achieve substantial changes in sphingolipid homeostasis and macrophage inflammatory response.

### Slide 10
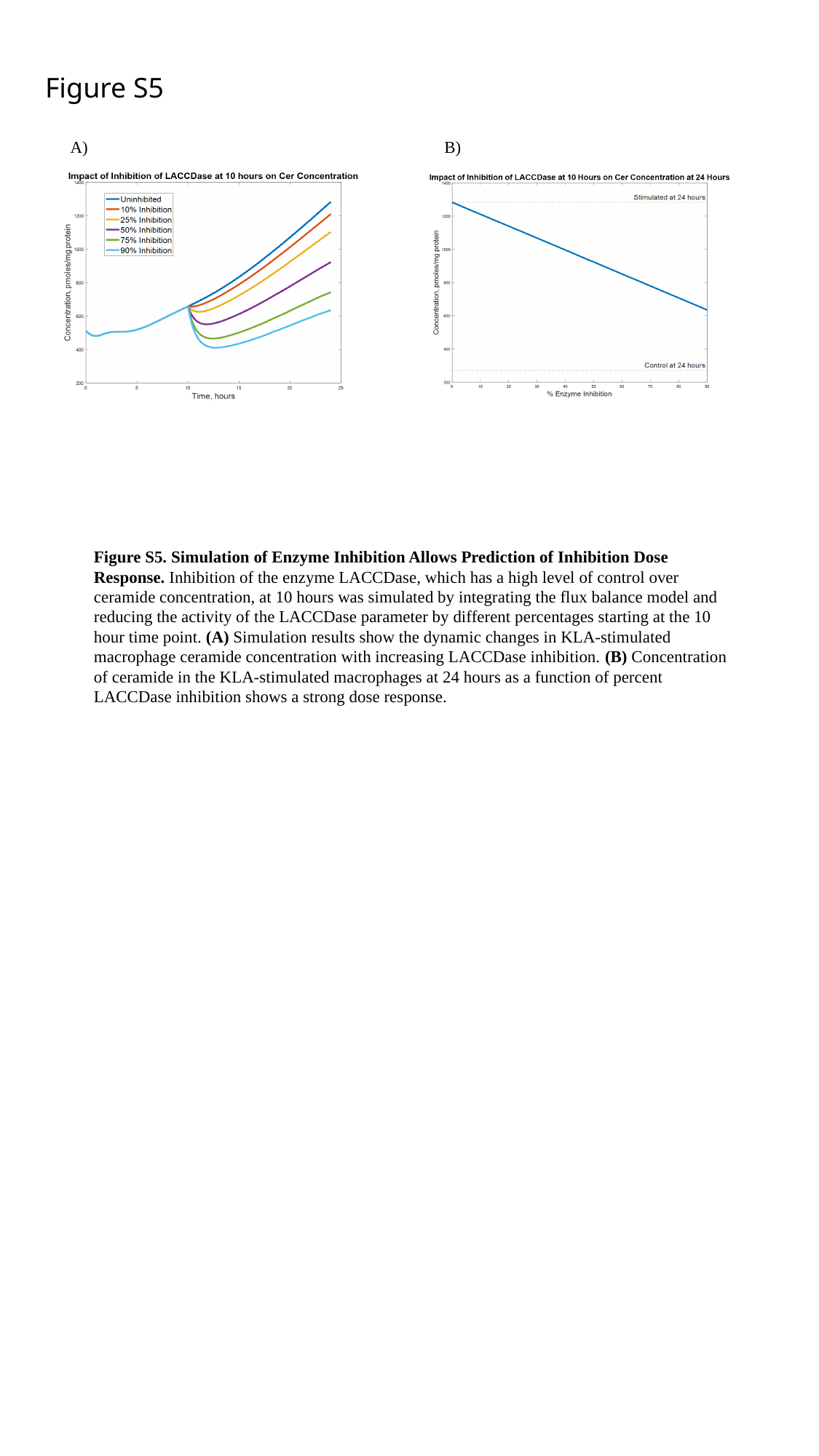

Figure S5
B)
A)
Figure S5. Simulation of Enzyme Inhibition Allows Prediction of Inhibition Dose Response. Inhibition of the enzyme LACCDase, which has a high level of control over ceramide concentration, at 10 hours was simulated by integrating the flux balance model and reducing the activity of the LACCDase parameter by different percentages starting at the 10 hour time point. (A) Simulation results show the dynamic changes in KLA-stimulated macrophage ceramide concentration with increasing LACCDase inhibition. (B) Concentration of ceramide in the KLA-stimulated macrophages at 24 hours as a function of percent LACCDase inhibition shows a strong dose response.

### Slide 11
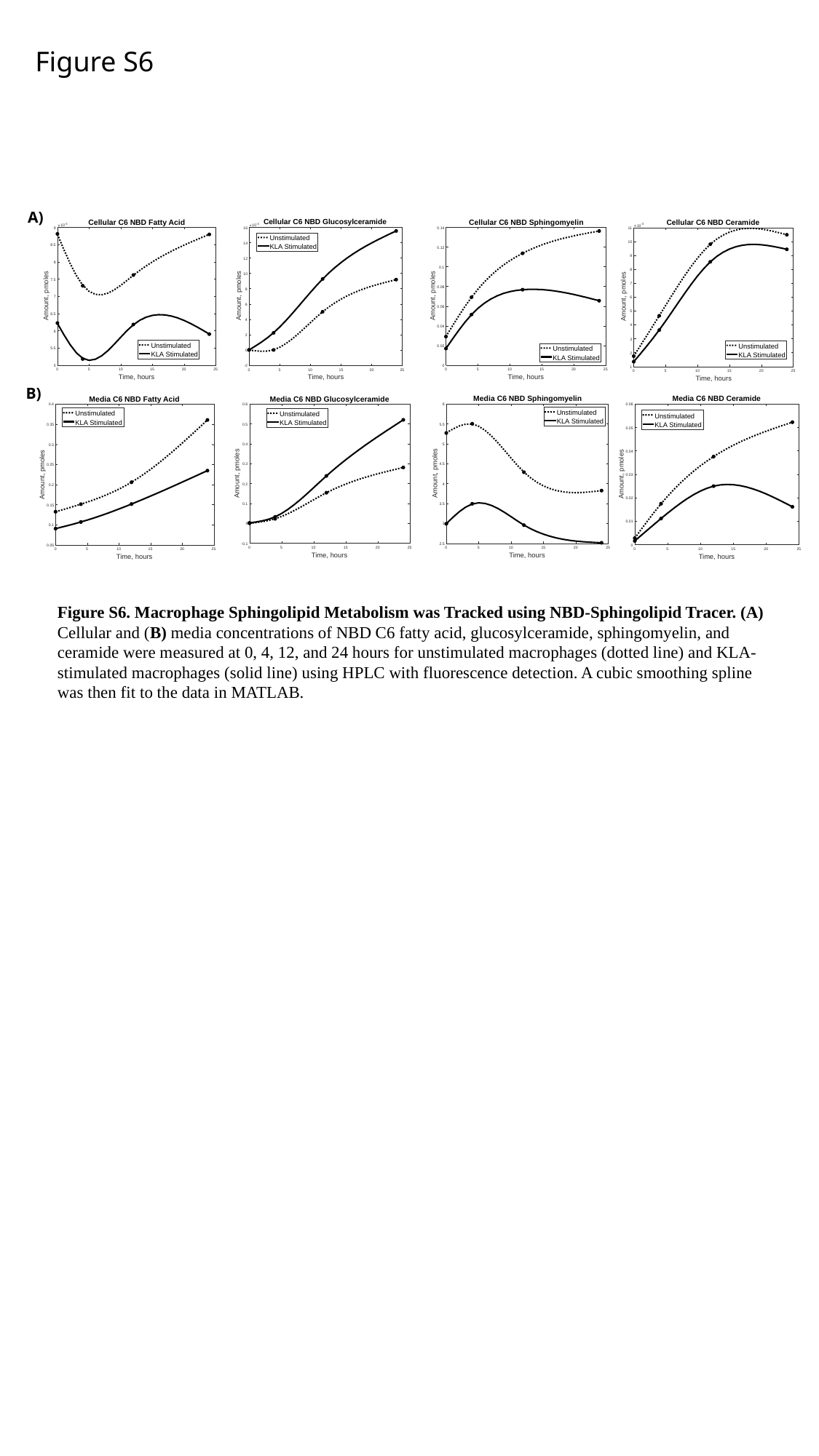

Figure S6
A)
B)
Figure S6. Macrophage Sphingolipid Metabolism was Tracked using NBD-Sphingolipid Tracer. (A) Cellular and (B) media concentrations of NBD C6 fatty acid, glucosylceramide, sphingomyelin, and ceramide were measured at 0, 4, 12, and 24 hours for unstimulated macrophages (dotted line) and KLA-stimulated macrophages (solid line) using HPLC with fluorescence detection. A cubic smoothing spline was then fit to the data in MATLAB.
